# MUNIn (<u>M</u>ultiple sample <u>UN</u>ifying long-range chromatin <u>In</u>teraction detector): a statistical framework for identifying long-range chromatin interactions from multiple samples

**DOI:** 10.1101/2020.11.12.380782

**Authors:** Weifang Liu, Armen Abnousi, Qian Zhang, Naoki Kubo, Joshua S. Martin Beem, Yun Li, Ming Hu, Yuchen Yang

## Abstract

Chromatin spatial organization (interactome) plays a critical role in genome function. Deep understanding of chromatin interactome can shed insights into transcriptional regulation mechanisms and human disease pathology. One essential task in the analysis of chromatin interactomic data is to identify long-range chromatin interactions. Existing approaches, such as HiCCUPS, FitHiC/FitHiC2 and FastHiC, are all designed for analyzing individual cell types or samples. None of them accounts for unbalanced sequencing depths and heterogeneity among multiple cell types or samples in a unified statistical framework. To fill in the gap, we have developed a novel statistical framework MUNIn (**<u>M</u>**ultiple sample **<u>UN</u>**ifying long-range chromatin **<u>In</u>**teraction detector) for identifying long-range chromatin interactions from multiple samples. MUNIn adopts a hierarchical hidden Markov random field (H-HMRF) model, in which the status (peak or background) of each interacting chromatin loci pair depends not only on the status of loci pairs in its neighborhood region, but also on the status of the same loci pair in other samples. To benchmark the performance of MUNIn, we performed comprehensive simulation studies and real data analysis, and showed that MUNIn can achieve much lower false positive rates for detecting sample-specific interactions (33.1 - 36.2%), and much enhanced statistical power for detecting shared peaks (up to 74.3%), compared to uni-sample analysis. Our data demonstrated that MUNIn is a useful tool for the integrative analysis of interactomic data from multiple samples.

## Introduction

Chromatin spatial organization plays a critical role in genome function associated with many important biological processes including transcription, DNA replication, and development.^1,2^ Recently, the ENCODE and the NIH Roadmap Epigenomics projects have identified millions of *cis*-regulatory elements (CREs; e.g., enhancers, silencers and insulators) in mammalian genomes. Notably, the majority of genes are not regulated by CREs in one-dimensional (1D) close vicinity. Instead, by forming three-dimensional (3D) long-range chromatin interactions, CREs are able to regulate the expression of genes hundreds of kilobases (kb) away. Deep understanding of chromatin interactome can shed light on gene regulation mechanisms, and reveal functionally causal genes underlying human complex diseases and traits. Comprehensive characterization of chromatin interactome has become an active research area since the development of Hi-C technology in 2009.^3^ Later on, Hi-C and other chromatin conformation capture (3C)-derived technologies (e.g., capture Hi-C, ChIA-PET, PLAC-Seq and HiChIP) have been widely used and great strides have been made to link chromatin interactome to mechanisms of transcriptional regulation and complex human diseases, including autoimmune diseases, neuropsychiatric disorders and cancers.^4–7^

Recent studies have shown that interactomes are highly dynamic across tissues, cell types, cell lines, experimental conditions, environmental triggers, and/or biological samples.^8,9^ Better characterization of such interactomic dynamics will substantially advance our understanding of transcription regulation across these conditions. To achieve this goal, one could use methods developed for single sample (for brevity, we use samples to denote multiple datasets across tissues, cell types, cell lines, experimental conditions, etc). However, such uni-sample analysis would fail to borrow information across samples, thus lose information for shared features, as well as resulting in false positives for sample-specific features. Presumably, as shown in eQTL analysis, shared (among at least two cell types) features typically contribute to a considerable proportion and increase with the number of cell types measured.^10^ For delineating shared and sample-specific features, Bayesian modeling has been shown repeatedly to boast the advantage of adaptively borrowing information such that little power loss incurs for sample-specific features while power to detect shared features increases substantially, as demonstrated in many genomic applications including gene expression, GWAS, ChIP-seq, population genetics, and microbiome.^11–15^

In this paper, we focus on the identification of statistically significant long-range chromatin interactions (“peaks” for short) from Hi-C data generated from multiple samples. The primary goal is the detection of both shared (i.e., shared by more than one samples) and sample-specific peaks. Existing Hi-C peak calling methods, such as HiCCUPS,^16^ FitHiC/FitHiC2^17,18^ and FastHiC,^19^ are all designed for calling peaks from single sample. None of them is able to account for unbalanced sequencing depths and heterogeneity among multiple samples in a unified statistical framework. To fill in the methodological gap, we propose MUNIn (**<u>M</u>**ultiple-sample **<u>un</u>**ifying long-range chromatin **<u>in</u>**teraction detector) for multiple samples Hi-C peak calling analysis. MUNIn adopts a hierarchical hidden Markov random field (H-HMRF) model, an extension of our previous HMRF peak caller.^20^ Specifically, in MUNIn, the status of each interacting chromatin loci pair (peak or background) depends not only on the status of loci pairs in its neighborhood region, but also on the status of the same loci pair in other closely related samples (Figure 1). Compared to uni-sample analysis, the H-HMRF approach adopted by MUNIn has the following three key advantages: (1) MUNIn can achieve lower false positive rates for the detection of sample-specific peaks. (2) MUNIn can achieve high power for the detection of shared peaks. (3) MUNIn can borrow information across all samples proportional to the corresponding sequencing depths. We have conducted comprehensive simulation studies and real data analysis to showcase the advantages of MUNIn over other Hi-C peak calling approaches.

**Figure 1.**
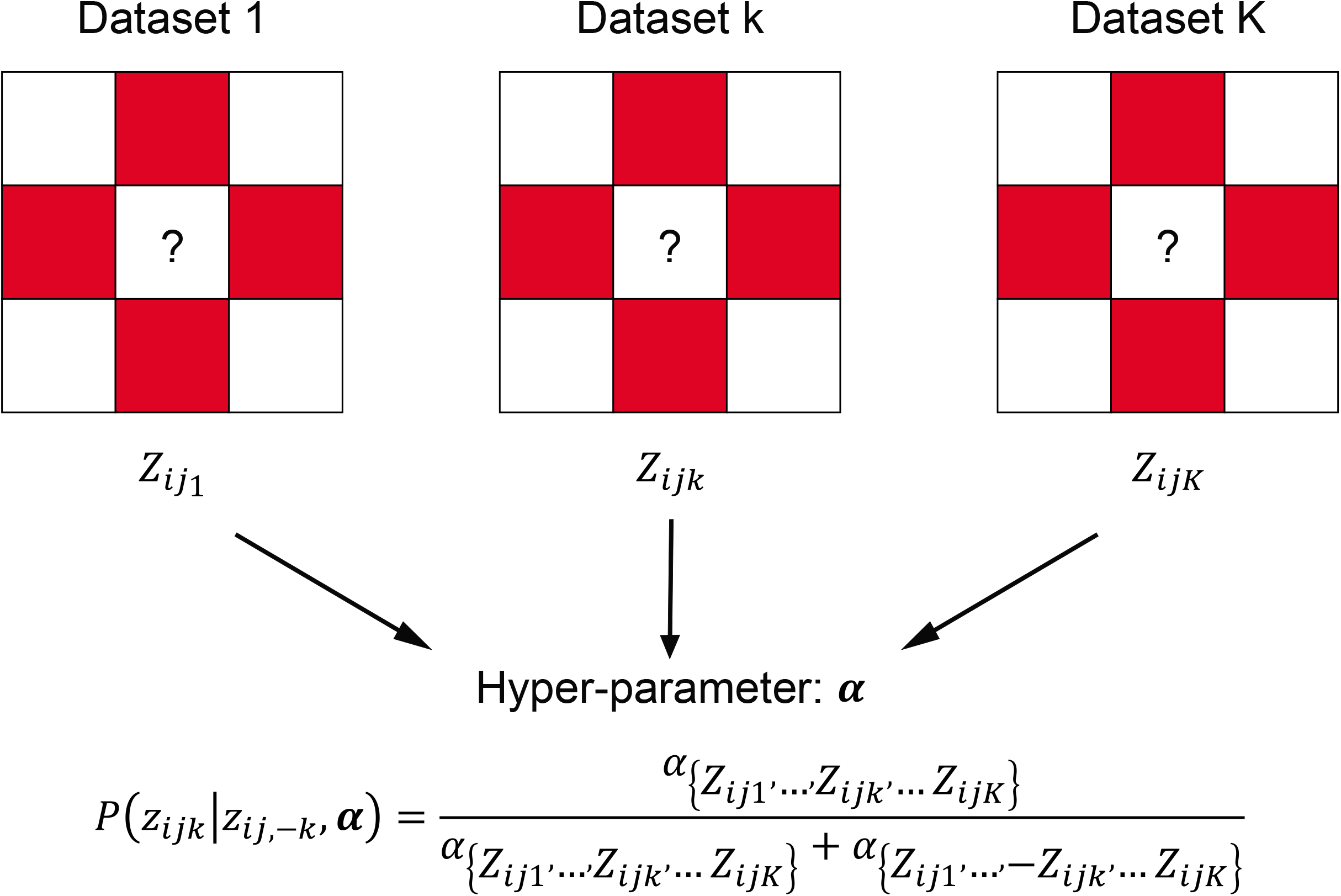
Statistical Schematics of MUNIn. In MUNIn, the chromatin interaction status (illustrated with question marks) of each loci pair (*i, j*) in a sample depends on not only the status of loci pairs in its neighborhood region (red blocks), but also the status of the same loci pair in other samples. Specifically, we model sample dependency by α, where the status of the (*i, j*)th pair in sample *k, z_ijk_*, depends on the status of the same (*i, j*)th pair in the other *K*-1 samples, given by the formula shown in the figure. Dependency on neighboring loci pairs captured by the hierarchical Ising prior. See Methods and Supplemental Section 1 for details.

## Materials and methods

### Overview of Statistical Modeling of MUNIn

Let *X_ijk_* and *e_jkk_* represent the observed and expected chromatin contact frequency spanning between bin *i* and bin *j* in sample (1 ≤ *i* < *j* ≤ *N*, 1 ≤ *k* ≤ *K*), respectively, where *N* is the total number of bins, and *K* is the total number of samples. *e_ijk_* is pre-calculated by FitHiC.^17^ Briefly, FitHiC used a non-parametric approach to estimate the empirical null distribution of contact frequency (detailed in **Supplemental Section 1**). We assume that *x_ijk_* follows a negative binomial (NB) distribution with mean *μ_ijk_* and overdispersion *ϕ_k_*:

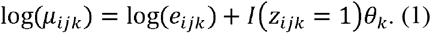

Here *Z_ijk_* ∈ {−1,1} is the peak indicator for bin pair (*i, j*), where *Z_ijk_* = 1 indicates (*i, j*) is a peak in sample *k* and *z_ijk_* = −1 otherwise. *θ_k_* is the signal-to-noise ratio in sample *k*. In other word, if (*i, j*) is a peak in sample *k, x_ijk_* follows the NB distribution *NB*(*e_ijk_* * exp{*θ_k_*}, *ϕ_k_*). If (*i, j*) is a background (i.e., non-peak) in sample *k, x_ijk_* follows the NB distribution *NB*(*e_ijk_, ϕ_k_*).

Then, we use a full Bayesian approach for statistical inference, and assign priors for all parameters (*z_ijk_,θ_k_, ϕ_k_*). Specifically, we adopt a hierarchical Ising prior to simultaneously model spatial dependency among *z_ijk_*’s within the same sample (i.e., for *z_ijk_*, borrowing information from *z_i′j′k_*:{|*i*′ – *i*| + |*j′* – *j* | = 1}), and the dependency across sample for the same pair (i.e., borrowing information from *z_ijk_′* with *k′* ∈ {1,…, *k* – 1, *k* + 1,…, *K*}). First of all, to model spatial dependency of peak indicator within sample *k*, we assume that:

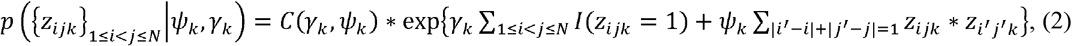

where *ψ_k_>* 0 is the inverse temperature parameter modeling the level the spatial dependency in sample *k, γ_k_* models the peak proportion in sample *k*, and *C* (*γ_k_, ψ_k_*) is the normalization constant. In addition, we model the heterogeneity of peak status for a given bin pair (*i, j*) among multiple samples, where the vector 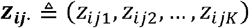 can take 2*^K^* possible configurations. We model them using a multinomial distribution 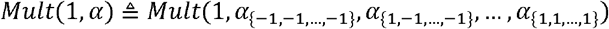. Here *α*_{−1, −1,…,−1}_ is the probability that the (*i, j*) pair is background in all *K* samples, *α*_{1, −1,…,−1}_ is the probability that the (*i, j*) pair is a peak in the first sample, but background in all the other *K* – 1 samples, and similarly *α*_{1,1,…,1}_ is the probability that the (*i, j*) pair is a peak in all *K* samples. Let *n_z_ij__* represent the frequency of a specific configuration *α_z_ij__*. The joint distribution is as follows:

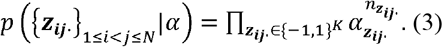

In this prior distribution, the peak probability of the (*i, j*) pair in sample *k* (*Z_ijk_*) depends on the status of same (*i, j*) pair in the other *K* – 1 samples:

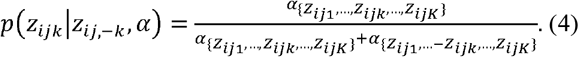

From the Bayes formulation, we have the joint posterior distribution as follows:

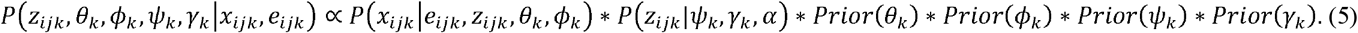

We used uniform prior distributions for *θ_k_,ϕ_k_,ψ_k_,γ_k_*, which were initialized from estimates from uni-sample analysis in our implementation (**Supplemental Section 2 and Section 3**). One key computational challenge is that in the proposed hierarchical Ising prior, the normalization constant involves *ψ_k_, γ_k_* and *α* is computationally prohibitive, since evaluating such normalization constant requires evaluating all 2^*K*N*(*N*−1)/2^ possible configurations of the peak indicator {*Z_ijk_*},. To address this challenge, we adopt a pseudo-likelihood approach, using the product of marginal likelihood to approximate the full joint likelihood. We have shown that such approximation leads to gains in both statistical and computational efficiency.^19^ Let {*Z*_−*i*,−*j,k*_} denote the set 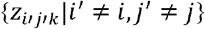 and {*Z_ij,−k_*} denote the set 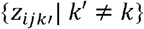, the posterior probability can be approximated by:

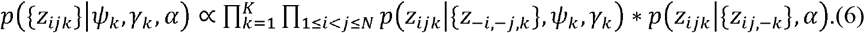

We use the Gibbs sampling algorithm to iteratively update each parameter. Details of statistical inference can be found in **Supplemental Section 2**.

### Simulation Framework

To benchmark the performance of MUNIn, we first performed simulation studies with three samples, where each sample represents a cell type, considering two scenarios: 1) all three samples had the same sequencing depth, and 2) the sequencing depth in sample 3 was half of that in sample 1 and sample 2. Each simulated sample consisted of a 100 × 100 contact matrix. To ensure the three samples were symmetric, we first simulated the peak status for one “hidden” sample using the Ising prior, where the two parameters *ψ_k_* was set to 0.2 and *γ_k_* was set to {0, −0.02, −0.05, −0.2, −0.4}, respectively. 10,000 Gibbs sampling steps were carried out to update peak status. Let: 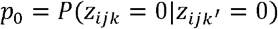 and 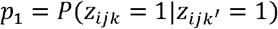 denote the level of dependence across samples. The peak status of the three testing samples were simulated from the hidden sample following three different sample-dependence levels *P*_0_ = *P*_1_ = 0.5, 0.8, or 0.9, where *P*_0_ = *P*_1_ = 0.5 indicates the peak status of three samples are independent, while *P*_0_ = *P*_1_ = 0.8 or 0.9 indicate the peak status of three samples are of median and high correlation. To simulate Hi-C data with equal sequencing depth, we specified expected contact frequency for the bin pair (*i, j*) to be inversely proportional to the genomic distance between two interacting anchor bins, following the same formula in each sample *k* (note the formula does not depend on *k*):

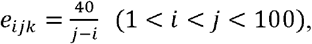

To simulate Hi-C data with different sequencing depths, we defined the expected count for bin pair (*i, j*) in sample 3 as:

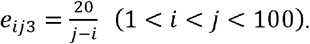

Next, we simulated the observed count from a negative binomial distribution:

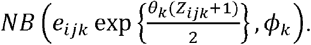

Here, the signal-to-noise ratio parameter *θ_k_* and the over-dispersion parameter *ϕ_k_* were set to be 1.5 and 10.0, respectively.

Simulations under each scenario were performed 100 times with different random seeds. We then applied both MUNIn and uni-sample analysis using single sample HMRF model (detailed in **Supplemental Section 3**) on simulated data of each scenario. The peak status was identified from the simulated data using both MUNIn and uni-sample method, and compared to the ground truth. ROC curve was computed using the *pROC* package.^21^ Furthermore, the performance of MUNIn was also evaluated according to the overall percentage of error in peak status *z_ijk_*, and the power and type I error for four types of peak status (i.e., shared, sample1-specific, sample2-specific and sample3-specific peaks), respectively.

### Performance evaluation

To evaluate the performance of MUNIn in real data, we first compared MUNIn to uni-sample analysis to two biological replicates of Hi-C data from human embryonic stem cells at 10kb resolution^22^ (**Table S1**), where the peak status is expected to be highly similar. For each biological replicate, both methods were implemented for peak calling within each topologically associating domain (TAD) of chromosome 1, where TADs were directly obtained from the original paper defined by the insulation score.^22^ To measure the consistency between these two replicates, we computed Adjusted Rand Index (ARI)^23^ for the peak status within each TAD.

Additionally, we also analyzed Hi-C data from two different cell lines, GM12878 and IMR90 at 10kb resolution^16^ (**Table S1**), again using both MUNIn and uni-sample analysis. Analyses were performed with each TAD in all chromosomes. Since some TAD boundaries are different between GM12878 and IMR90, we first defined the overlapped TAD regions as the shared TADs between two samples, and only retained the shared TADs spanning at least 200kb for the downstream analysis. Sample dependency was inferred for each TAD based on the results of uni-sample analysis. Since there is no ground truth for peaks, we selected significant chromatin interactions (p-value < 0.01 and raw interaction frequency > 5) identified by promoter-capture Hi-C (PC-HiC)^9^ in GM12878 and IMR90 cells as the working truth (**Table S1**). Since significant interactions identified from PC-HiC data are enriched of promoters, we filtered our significant peaks to only remain bin pairs where at least one of two bins overlaps with a promoter. The detailed evaluation framework is in **Supplemental Section 4**. We did additional performance evaluation by running MUNIn by a sliding window approach instead of shared TADs, and we also performed peak calling on samples under different conditions from mouse embryonic stem cells for both wild-type (without CTCF depletion) and after CTCF deletion resolution^26^ (**Table S1**; **Supplemental Section 5**).

## Results

### Simulation results

To evaluate the performance of MUNIn, we first conducted simulation studies with three samples, considering two scenarios: (1) all three samples have equal sequencing depth, and (2) the sequencing depth in sample 3 is half of that in sample 1 and 2. In both scenarios, MUNIn outperforms uni-sample analysis (**Figures 2–3; Figures S1-S4**). In the first scenario, when all three samples are independent (*p*_0_ = *p*_1_ = 0-5), MUNIn achieves comparable results to uni-sample analysis, where the medians of the overall error rate (denoted as “%error”) in peak identification of MUNIn range from 16.3 to 16.4% and those of uni-sample analysis are 17.2 - 17.3% (**Figure 2a**). With increased sample dependency, MUNIn achieves lower %error than uni-sample analysis. When the sample dependency becomes high, MUNIn reduces %error by approximately 30.3% on top of uni-sample results (11.9 - 12.0% for MUNIn and 17.0 - 17.2% for uni-sample analysis) (**Figure 2a**). We then assessed the power and type I error for detecting shared and sample-specific peaks by MUNIn and uni-sample analysis. When three samples are highly correlated, MUNIn achieves substantial power gain in shared peaks across samples than uni-sample analysis (85.9% vs. 54.1%; **Figure 2c**), at the cost of a slight increase in error rate (20.6% vs. 9. 1%; **Figure 2d**). In addition, MUNIn reduces the type I error in calling sample-specific peaks by 33.1 - 34.3% on the top of uni-sample results (45.5 - 46.3% vs. 69.3 - 69.5%; **Figure S1a**), at the cost of power loss (36.4 - 37.1% vs. 57.3 - 58.5%; **Figure S1b**). The ROC curves showed that MUNIn better detects shared peaks than uni-sample analysis (**Figure 2b**), and these two methods performed comparably in sample-specific peaks (**Figure S2**).

**Figure 2.**
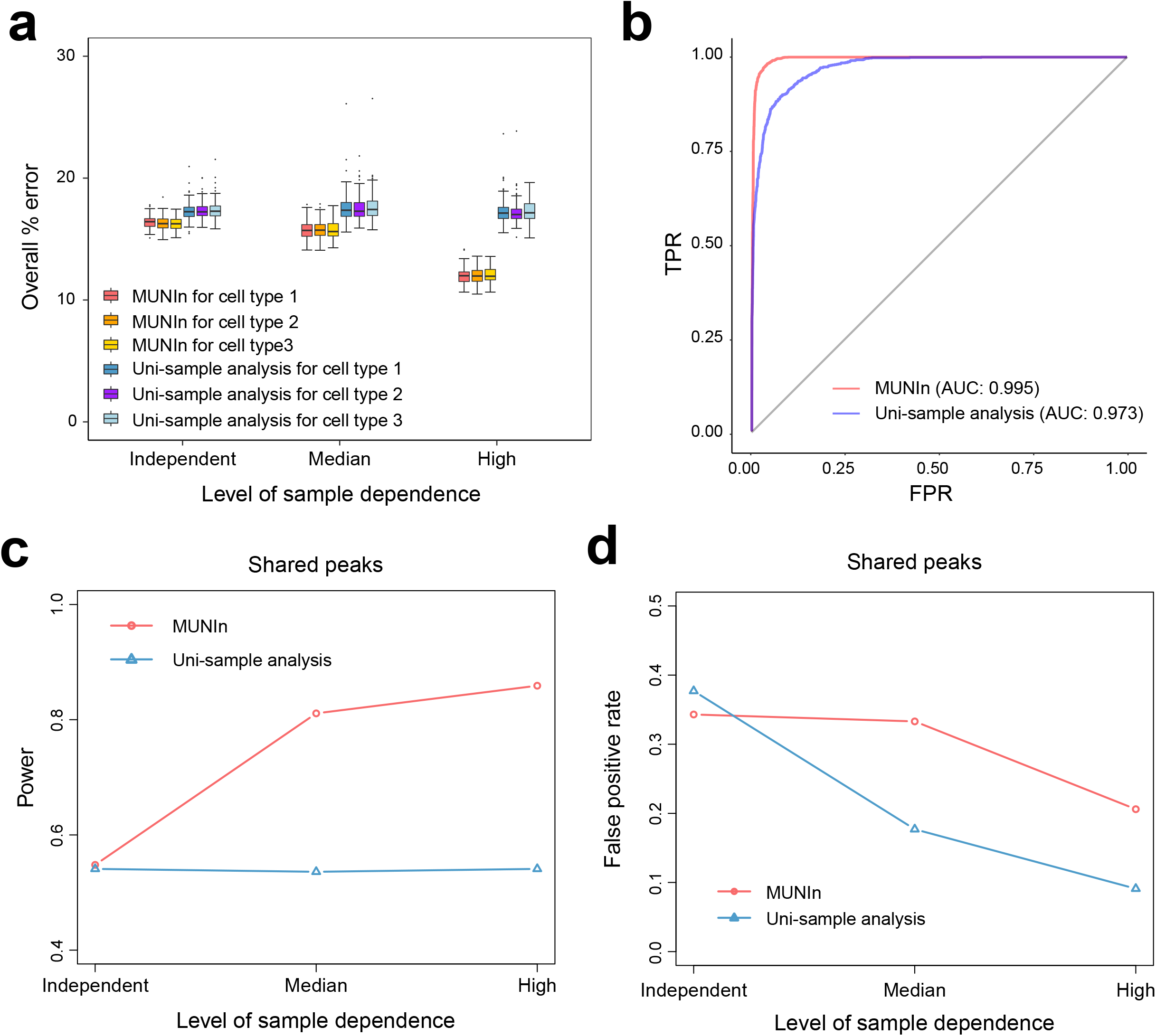
Performance comparison between MUNIn and uni-sample analysis in the simulation data where all three samples have equal sequencing depth. (**a**) The overall error rate (denoted as “%error”) in peak identification in each sample using MUNIn and unisample analysis. (**b**) ROC curves for shared peaks identified by MUNIn and uni-sample analysis. (**c**) Power for the shared peaks identified using MUNIn and uni-sample analysis. (**d**) False positive rate for the shared peaks identified by MUNIn and uni-sample analysis.

**Figure 3.**
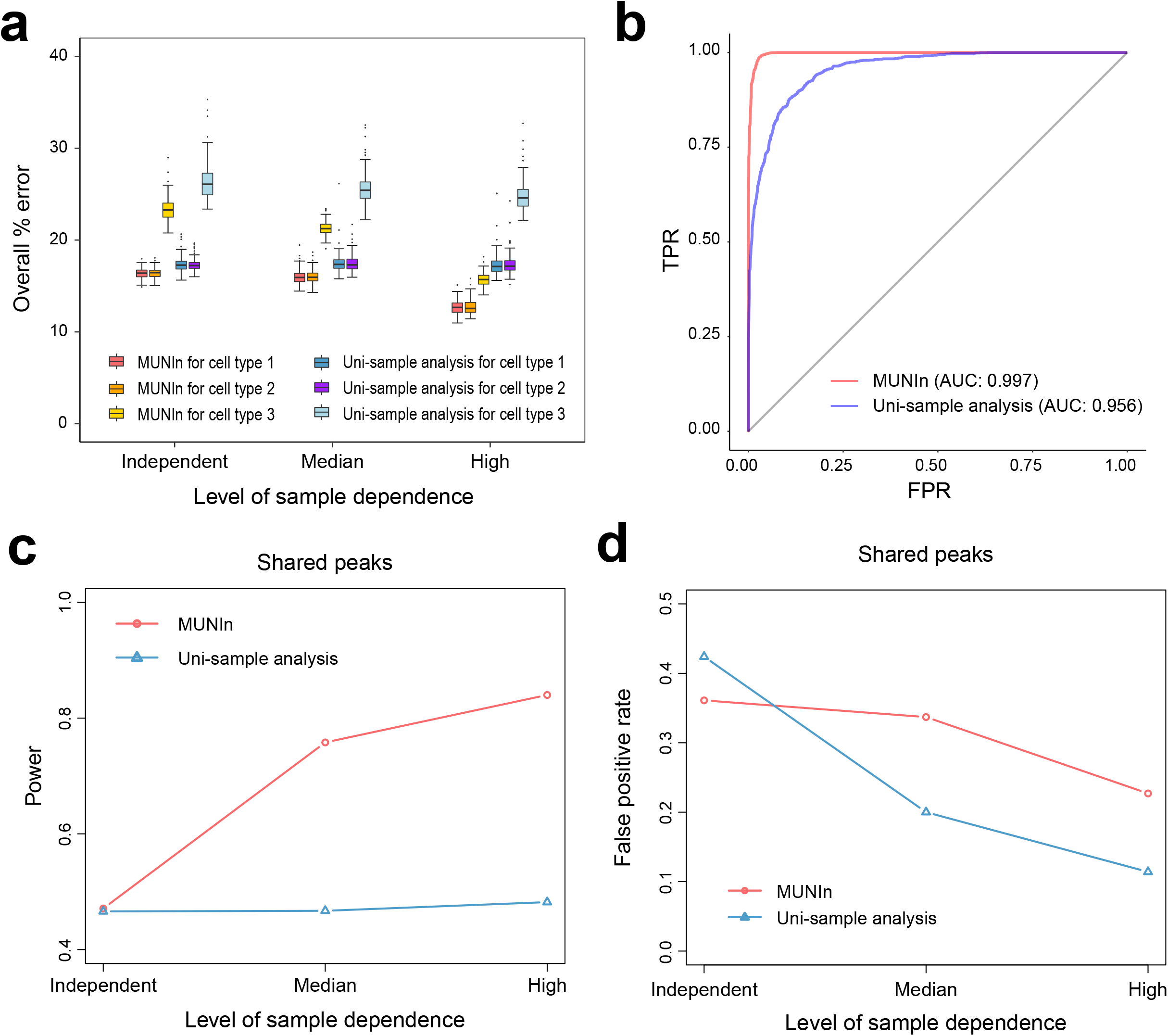
Performance comparison between MUNIn and uni-sample analysis in the simulation data where the sequencing depth in sample 3 is half of that in sample 1 and 2. (**a**) The overall error rate (denoted as “%error”) in peak identification in each sample using MUNIn and uni-sample analysis. (**b**) ROC curves for shared peaks identified by MUNIn and uni-sample analysis. (**c**) Power for the shared peaks identified using MUNIn and uni-sample analysis. (**d**) False positive rate for the shared peaks identified by MUNIn and uni-sample analysis.

Furthermore, when three samples are with different sequencing depths, we observe consistent patterns that MUNIn outperforms uni-sample analysis, especially for sample 3 with shallower sequencing depth (**Figure 3; Figures S3-S4**). Similar to scenario 1, the ROC curves show that MUNIn exhibits better calling in shared peaks (**Figure 3b**). Consistently, MUNIn substantially improves the power in calling shared peaks than uni-sample analysis (84.0% vs 48.2% by MUNIn and uni-sample analysis, respectively) with a slightly increase of type I error (22.7% vs 11.4%) (**Figure 3c; Figure 3d**). More importantly, MUNIn achieves 36.2% reduction of %error for sample 3 with shallower sequencing depth on the top of uni-sample analysis results with high sample dependence (15.7% vs 24.6%; **Figure 3a**). MUNIn also attains lower type I error in calling sample3-specific peaks (51.1% vs 74.4%) with a loss in power (26.7% vs 48.1%) (**Figure S3a; Figure S3b**). These results indicate that MUNIn can accurately identify peaks in the shallowly sequenced sample by adaptively borrowing information from deeply sequenced samples. We further evaluated the robustness and scalability of MUNIn using simulation data where we evaluated results with non-zeros *γ_k_s* and increased sample size (**Supplemental Section 5**; **Figure S5 and S6**).

### Real data analysis

To assess the performance of MUNIn in real data, we compared the consistency of peaks status between two replicates of human embryonic stem cells between MUNIn and uni-sample analysis. Comparatively, the ARI values of MUNIn are significantly higher than those of uni-sample analysis (Wilcoxon test, *p*-value < 2.2e-16; **Figure 4**; **Figure S7**). Specifically, the median value of ARI in MUNIn is 0.993, which shows 48.9% improvement over that of uni-sample analysis (**Figure S7**). Our results suggest improved consistency between two replicates by MUNIn, compared to uni-sample analysis.

**Figure 4.**
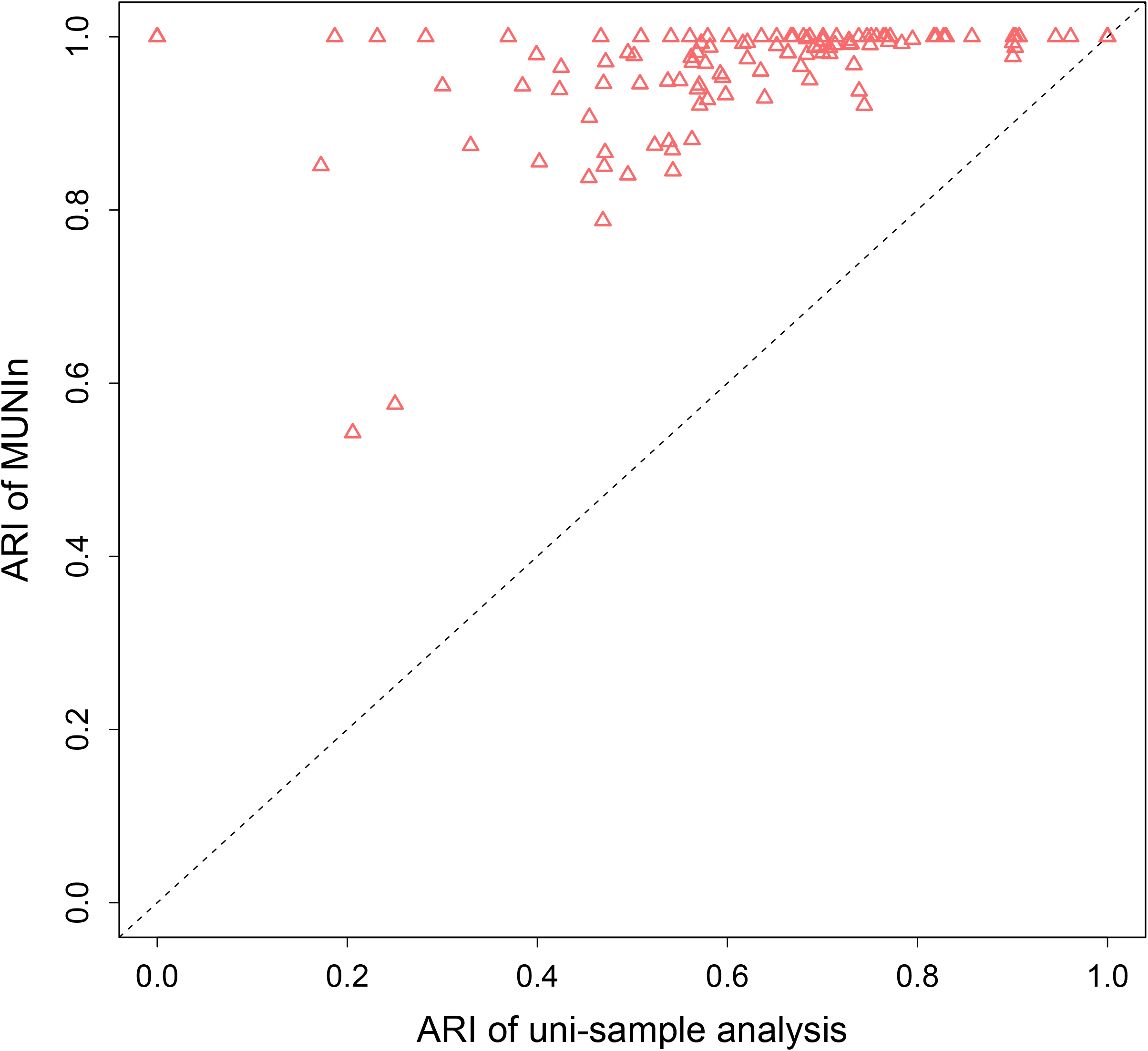
Adjusted Rand Index (ARI) showing the consistency of peak calling by MUNIn and uni-sample analysis between the two replicates of human embryonic stem cells. Each triangle represents a TAD. The x- and y-axis: ARI of uni-sample analysis and MUNIn, respectively.

We further compared the accuracy of peak calling in GM12878 and IMR90 cell lines between MUNIn and uni-sample analysis. In total, 439,412 and 432,394 shared peaks were detected by MUNIn and uni-sample analysis, respectively, 376,658 of which were shared by both methods (85.7 and 87.1% of the shared peaks identified MUNIn and uni-sample analysis, respectively) (**Figure S8a**). 217,400 and 82,614 GM12878 and IMR90-specific peaks were identified by MUNIn, while 315,849 and 141,708 GM12878 and IMR90-specific peaks were detected by uni-sample analysis. Among them, 77.5 and 75.7% of GM12878- and IMR90-specfic peaks called by MUNIn were also identified by uni-sample analysis (**Figure S8b and c**). The ROC curves show that MUNIn obtains more accurate results for both GM12878 and IMR90-specific peaks (**Figure 5a and d**), while its performance in shared peaks is comparable to uni-sample analysis (**Figure S9**). The area under the curve (AUC) for GM12878 and IMR90-specific peaks of MUNIn increases by 3.0% and 4.5%, respectively, on top of uni-sample analysis (**Figure 5a and d**). One example of GM12878-specific peak exclusively identified by MUNIn is shown in **Figure 5b (Figure S10).** One bin of this pair is overlapped with the promoter of *ZNF827* gene (transcription start site (TSS) +/− 500 bp), while the other is overlapped with a known typical enhancer in GM12878 cells (**Figure S11**).^24^ In addition, *ZNF827* showed higher gene expression in GM12878 cells than in IMR90 cells (**Figure 5c**), which further suggests the potential role of this GM12878-specific peak in cell-type-specific transcriptional regulation gene. Similarly, the MUNIn-exclusively identified peak between bins chr4:95,000,000-95,010,000 and chr4:95,170,000-95,180,000 is specific to IMR90, which is involved in the regulation of *F3* gene (**Figure 5e; Figure S12**). *F3* gene encodes the tissue factor coagulation factor III and it is usually expressed in the fibroblasts surrounding blood vessels. Consistently, we observed a higher expression level of *F3* gene in IMR90 cells than in GM12878 cells (**Figure 5f**). Additional real data evaluation also showed value of borrowing information across samples where we compared MUNIn to uni-sample analysis and FitHiC (**Supplemental Section 5**; **Figure S13-S17**).

**Figure 5.**
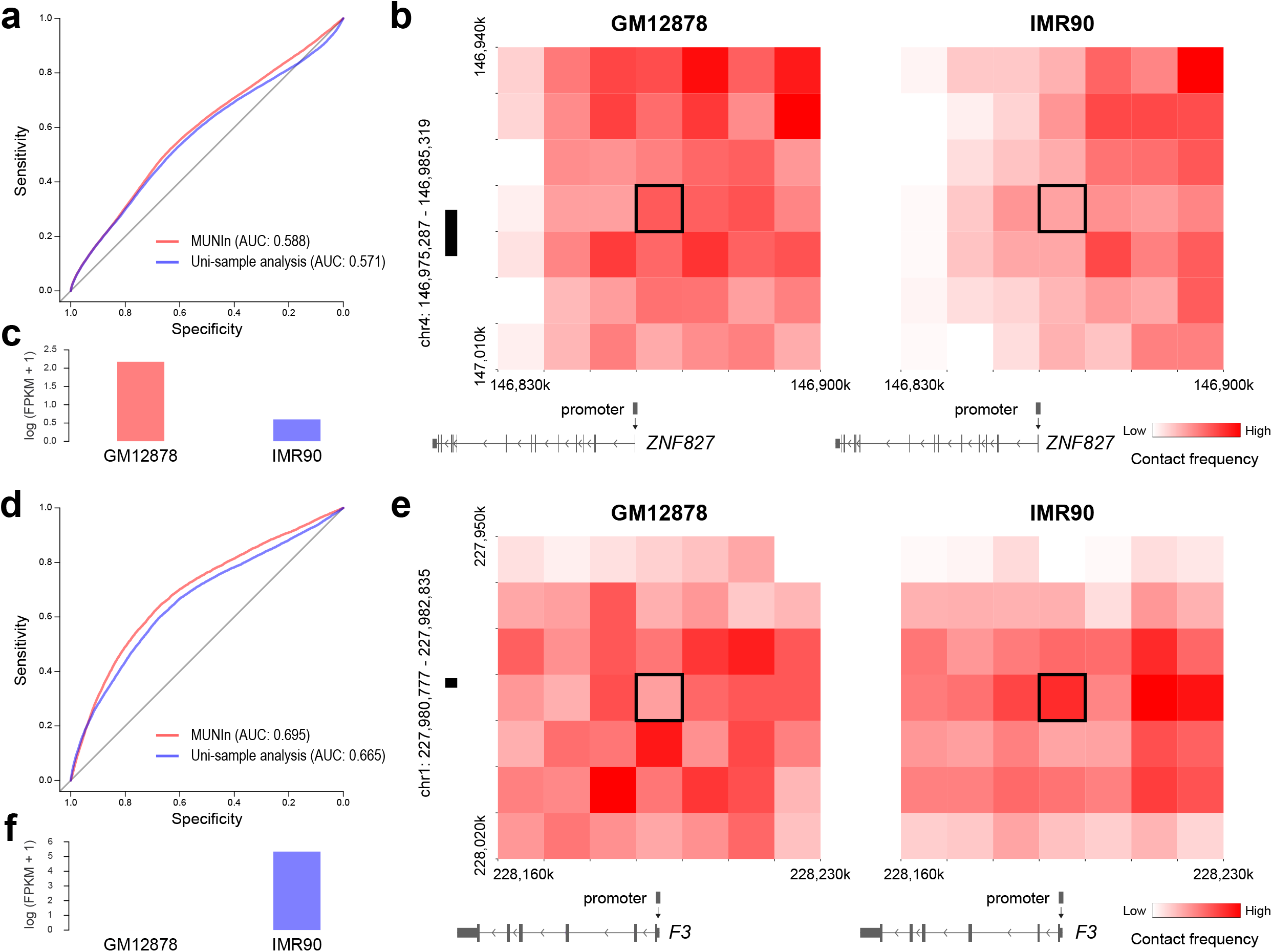
Performance comparison between MUNIn and uni-sample analysis in the Hi-C data of GM12878 and IMR90 cell lines. (**a**) ROC for GM12878-specfic peaks identified by MUNIn and uni-sample analysis. (**b**) Heatmap showing one example of the GM12878-specific peaks in GM12878 (left) and IMR90 (right) Hi-C data. One bin of this pair (highlighted in black) is overlapped with the promoter of *ZNF827* gene (transcription start site (TSS) +/− 500 bp), while the other is overlapped with a known typical enhancer (chr4:146,975,287-146,985,319) in GM12878 cells. Gene model is obtained from WashU epigenome browser.^25^ (**c**) Gene expression profiles of *ZNF827* in GM12878 and IMR90 cells. (**d**) ROC for IMR90-specfic peaks identified by MUNIn and uni-sample analysis. (**e**) Heatmap showing one example of the IMR90-specific peaks in GM12878 (left) and IMR90 (right) Hi-C data. One bin of this pair (highlighted in black) is overlapped with the promoter of *F3* gene, while the other is overlapped with a known typical enhancer (chr1:227,980,777-227,982,835) in IMR90 cells. Gene mode is obtained from WashU epigenome browser. (**f**) Gene expression profiles of *F3* gene in GM12878 and IMR90 cell lines.

## Discussion

In this study, we present MUNIn, a statistical framework to identify long-range chromatin interactions for Hi-C data from multiple tissues, cell lines or cell types. MUNIn is built on our previously developed methods, HMRF peak caller and FastHiC.^19,20^ On top of HMRF, MUNIn jointly models multiple samples and explicitly accounts for the dependency across samples. It simultaneously accounts for both spatial dependency within each sample, and dependency across samples. By adaptively borrowing information in both aspects, MUNIn can enhance the power of detecting shared peaks, and reduce type I error of detecting sample-specific peaks.

MUNIn exhibits substantial advantages in calling peaks shared across samples compared to uni-sample analysis (**Figure 2b**), which are more pronounced with the increased level of across sample dependency. In addition, with imbalanced sequencing depth among different samples, uni-sample analysis may mis-classify shared peaks as sample-specific due to differential power across samples. Comparatively, MUNIn can more accurately identify shared peaks (**Figure 3b**). Noticeably, MUNIn resulted in reduced false positives when calling sample-specific peaks for the sample with shallower depth (**Figure S3a**). This is because MUNIn can borrow information from samples with higher sequencing depth based on the level of dependency across samples, which is also learned from the data. In our real data evaluations, MUNIn also outperformed uni-sample analysis. Specifically, for Hi-C data from human embryonic stem cells, MUNIn exhibited significantly higher consistency between the two biological replicates than the uni-sample analysis (**Figure 4; Figure S7**). For Hi-C data from GM12878 and IMR90 cell lines, MUNIn more accurately identified cell-line specific peaks, in terms of both sensitivity and specificity (**Figure 5a and d**). In addition, GM12878- and IMR90-specific peaks exclusively identified by MUNIn showed in **Figure 5** may play a potential role in regulating gene *ZNF827* and gene *F3* respectively, which are differentially expressed between these two cell lines in the expected direction (**Figure 5c and f**). In our real data analysis, we ran MUNIn in shared TADs across samples instead of the whole chromosomes. We realized that regions outside of TADs or TADs that are not shared across samples may contain sample specific peaks, therefore we re-ran the analysis including those regions by a sliding window approach (**Figure S13**; **Supplemental Section 5**). Our results suggested that including those regions did not have a significant impact of the performance of MUNIn (**Figure S13**). Additionally, we assessed MUNIn’s performance on the Hi-C datasets from mouse embryonic stem cells for both wild-type (without CTCF depletion) and after CTCF deletion at 10kb resolution^26^ (**Table S1**). The results showed that MUNIn better captured the wild-type-specific pattern in mESC Hi-C data than uni-sample analysis and FitHiC (**Figure S14 and S15**; **Supplemental Section 5**), demonstrating the power of MUNIn to reveal peaks more powerfully and accurately by borrowing information from another sample.

Taking the advantages of jointly modeling multiple samples, MUNIN can easily accommodate many more samples simultaneously. MUNIn shows a high computational efficiency that MUNIn takes ~36 minutes to perform peak calling in a 2MB TAD of 10kb resolution (**Figure S16 and S17**; **Supplemental Section 5**). Moreover, MUNIn is also able to handle multiple samples with differential levels of dependency, for example, when samples forming clusters where samples within a cluster are more correlated than those across clusters. The MUNIn framework can be further extended to accommodate time series chromatin conformation data, which will be explored in our future work. Although MUNIn simultaneously models multiple samples, we note that the goal is to detect chromatin interactions of various peak status configurations across samples, rather than differential interactions. Theoretically, while the posterior probabilities of the peak status configurations can inform differential interactions, it is not our objective here and can be a direction for further exploration.

Taken together, our results show the advantages of MUNIn over the uni-sample approach when analyzing data from multiple samples. By adaptively borrowing information both within and across samples, MUNIn can achieve much improved power in detecting shared peaks, and much reduced type I error in detecting sample-specific peaks. MUNIn’s ability to reduce false positive sample-specific peak calls due to imbalanced sequencing depths across samples is also appealing. Finally, MUNIn can more effectively identify biologically relevant chromatin interactions with better sensitivity than the uni-sample strategy. We anticipate that MUNIn will become a convenient and essential tool in the analysis of multi-sample chromatin spatial organization data.

## Data and code availability

MUNIn is compiled as a C++ program, and freely available at https://github.com/yycunc/MUNIn and https://yunliweb.its.unc.edu/MUNIn/.

## Supplemental Data

Supplemental materials are available online.

## Acknowledgements

This research was supported by the National Institute of Health grants [R01 HL129132, U01 DA052713, R01 GM105785, and P50 HD103573].

## Web resources

WashU epigenome browser, http://epigenomegateway.wustl.edu/browser/

HUGIn, https://yunliweb.its.unc.edu/hugin/

