## Supplementary file for "MUNIn (<u>M</u>ultiple sample <u>UN</u>ifying long-range chromatin <u>In</u>teraction detector): a statistical framework for identifying long-range chromatin interactions from multiple samples"

**Supplemental Material**

**Section 1: Details of inferred expected contacts from FitHiC**

FitHiC models the expected contact count conditioned on observed contact counts and the genomic distance between interacting regions.^1^ Specifically, all locus pairs with a non-zero contact count are sorted with respect to increasing genomic distance between the two ends of the pair and this sorted list is broken into *b* bins. Then, the total number of contact counts is divided into the *b* bins using an equal occupancy binning strategy. Thus, all bins have approximately equal number of contacts. In each bin, the average genomic distance (x-axis) and contact probability (y-axis) are computed among all pairs (including possible pairs with zero contact counts) in this bin. Then, FitHiC fits a cubic smoothing spline (third degree polynomial) to these x, y values (one per bin) to learn a continuous function that relates these two entities. The inferred expected contact probabilities are further corrected through the bias values, which are computed per locus/bin by KR normalization method.^2^

**Section 2: Details of statistical inference**

Based on Bayes formula, we have the joint posterior distribution as follows:

$$P\left( \{z_{ijk}\},\{\theta_{k}\},{\{\phi}_{k}\},\{\psi_{k}\},\{\gamma_{k}\} | \{x_{ijk}\},{\{e}_{ijk}\} \right)\propto P\left( {\{x}_{ijk}\} | {\{e}_{ijk}\},{\{z}_{ijk}\},{\{\theta}_{k}\},\{\phi_{k}\} \right)*Prior\left( \{z_{ijk}\}|{\{\psi}_{k}\},{\{\gamma}_{k}\},\boldsymbol{\alpha} \right)*Prior\left( \{\theta_{k}\} \right)*Prior\left( \{\phi_{k}\} \right)*Prior\left( \{\psi_{k}\} \right)*Prior\left( \{\gamma_{k}\} \right)$$

Note that we used uniform prior distributions for $\theta_{k}{, \phi}_{k},\psi_{k},\gamma_{k}$, which were initialized from estimates from uni-sample analysis in our implementation (see **Section 3**).

Let $NB(x|\mu,\phi)$ represent the probability mass function of negative binomial distribution with mean $\mu$ and over-dispersion $\phi$, then

$$P\left( {\{x}_{ijk}\} | {\{e}_{ijk}\},\{z_{ijk}\},{\{\theta}_{k}\},{\{\phi}_{k}\} \right)=\prod_{k=1}^{K} \prod_{1\leq i<j\leq N} NB\left( x_{ijk} | e_{ijk}\exp\{I\left( z_{ijk}=1 \right)\theta_{k}\},\phi_{k} \right).$$

In addition, we use pseudo-likelihood approximation to calculate the hierarchical Ising prior where the joint density is approximated by the product of conditional densities:

$$P\left( {\{z}_{ijk}\}|{\{\psi}_{k}\},{\{\gamma}_{k}\},\boldsymbol{\alpha} \right)\approx\prod_{k=1}^{K} \prod_{1\leq i<j\leq N} P\left( z_{ijk} | {\{z}_{-\left( ijk \right)}\},\psi_{k},\gamma_{k},\boldsymbol{\alpha} \right).$$

Here

$$P\left( z_{ijk} | {\{z}_{-\left( ijk \right)}\},\psi_{k},\gamma_{k},\boldsymbol{\alpha} \right)\propto P\left( z_{ijk} | {\{z}_{-i,-j,k}\},\psi_{k},\gamma_{k} \right)*P\left( z_{ijk} | {\{z}_{ij,-k}\},\boldsymbol{\alpha} \right)\propto\exp\left\{ \gamma_{k}{I(z}_{ijk}=1)+\psi_{k}*z_{ijk}*\sum_{\left| i^{'}-i \right|+\left| j^{'}-j \right|=1} z_{i^{'}j^{'}k} \right\}*\frac{\alpha_{\{Z_{ij1},\ldots,Z_{ijk},\ldots,Z_{ijK}\}}}{\alpha_{\{Z_{ij1},\ldots,Z_{ijk},\ldots,Z_{ijK}\}}+\alpha_{\{Z_{ij1},\ldots,-Z_{ijk},\ldots,Z_{ijK}\}}}\propto\exp\left\{ \gamma_{k}{I(z}_{ijk}=1)+\psi_{k}*z_{ijk}*\sum_{|i^{'}-i|+|j^{'}-j|=1} z_{i^{'}j^{'}k} \right\}*\alpha_{\left\{ z_{ij1},\ldots z_{ijk},\ldots,z_{ijK} \right\}}.$$

Where $\{z_{-\left( ijk \right)}\}$ denotes the set {$z_{i'j'k'}|i^{'}\neq i, j^{'}\neq j, k'\neq k$}, $\{z_{-i,-j,k}\}$ denotes the set {$z_{i'j'k}|i^{'}\neq i, j^{'}\neq j$}, and $\{z_{ij,-k}\}$ denotes the set {$z_{ijk'}| k'\neq k$}.

Define:

$$A\left( z_{ijk} \right)=\exp\left\{ \gamma_{k}{I(z}_{ijk}=1)+\psi_{k}*z_{ijk}*\sum_{|i^{'}-i|+|j^{'}-j|=1} z_{i^{'}j^{'}k} \right\}*\alpha_{\left\{ z_{ij1},\ldots z_{ijk},\ldots,z_{ijK} \right\}}.$$

We have:

$$P\left( z_{ijk} | {\{z}_{-\left( ijk \right)}\},\psi_{k},\gamma_{k},\boldsymbol{\alpha} \right)=\frac{A\left( z_{ijk} \right)}{A\left( z_{ijk} \right)+A\left( -z_{ijk} \right)}=\frac{1}{1+A\left( -z_{ijk} \right)/A\left( z_{ijk} \right)}.$$

Taken together,

$$P\left( {\{z}_{ijk}\}|{{\{z}_{-\left( ijk \right)}\},\{\psi}_{k}\},\{\gamma_{k}\},\boldsymbol{\alpha} \right)=\prod_{k=1}^{K} \prod_{\begin{aligned} 1\leq i<j\leq N \\ \end{aligned}} \frac{1}{1+A\left( -z_{ijk} \right)/A\left( z_{ijk} \right)}.$$

The log of the joint posterior distribution is approximated as follows:

$$\log P\left( {\{z}_{ijk}\},{\{\theta}_{k}\},\{\phi_{k}\},\{\psi_{k}\},\{\gamma_{k}\} | {\{x}_{ijk}\},{\{e}_{ijk}\} \right)\approx Constant+ \sum_{k=1}^{K} \sum_{1\leq i<j\leq N} \log NB\left( x_{ijk} | e_{ijk}\exp\left\{ \frac{z_{ijk}+1}{2}\theta_{k} \right\},\phi_{k} \right)$$

$$+\log P\left( {\{z}_{ijk}\}|{{\{z}_{-\left( ijk \right)}\},\{\psi}_{k}\},\{\gamma_{k}\},\boldsymbol{\alpha} \right)$$

$$=Constant+\sum_{k=1}^{K} \sum_{1\leq i<j\leq N} \left\{ \log\Gamma\left( x_{ijk}+\phi_{k} \right)-\log\Gamma\left( \phi_{k} \right)+x_{ijk}\left( \log e_{ijk}+\theta_{k}\frac{z_{ijk}+1}{2} \right)+\phi_{k}\log\phi_{k}-\left( x_{ijk}+\phi_{k} \right)\log\left( e_{ijk}\exp\left\{ \theta_{k}\frac{z_{ijk}+1}{2} \right\}+\phi_{k} \right) \right\}-\sum_{k=1}^{K} \sum_{1\leq i<j\leq N} \log\left\{ 1+\frac{A\left( -z_{ijk} \right)}{A\left( z_{ijk} \right)} \right\}$$

In the equation above,

$$\frac{A\left( -z_{ijk} \right)}{A\left( z_{ijk} \right)}=\frac{\exp\left\{ -\gamma_{k}z_{ijk}-\psi_{k}*z_{ijk}*\sum_{\left| i^{'}-i \right|+\left| j^{'}-j \right|=1} z_{i^{'}j^{'}k} \right\}*\alpha_{\left\{ z_{ij1},\ldots{-z}_{ijk},\ldots,z_{ijK} \right\}}}{\exp\left\{ \gamma_{k}z_{ijk}+\psi_{k}*z_{ijk}*\sum_{\left| i^{'}-i \right|+\left| j^{'}-j \right|=1} z_{i^{'}j^{'}k} \right\}*\alpha_{\left\{ z_{ij1},\ldots z_{ijk},\ldots,z_{ijK} \right\}}}$$

$$=\exp\left\{ -2\gamma_{k}z_{ijk}-2\psi_{k}*z_{ijk}*\sum_{|i^{'}-i|+|j^{'}-j|=1} z_{i^{'}j^{'}k} \right\}*\frac{\alpha_{\left\{ z_{ij1},\ldots-z_{ijk},\ldots,z_{ijK} \right\}}}{\alpha_{\left\{ z_{ij1},\ldots z_{ijk},\ldots,z_{ijK} \right\}}}.$$

In the Gibbs sampler, the conditional distribution $z_{ijk}\in\{-1,1\}$ follows a Bernoulli distribution. We have:

$$\log P\left( z_{ijk}=1 | \theta_{k},\phi_{k},\psi_{k},\gamma_{k},x_{ijk},e_{ijk} \right)=x_{ijk}\theta_{k}-\left( x_{ijk}+\phi_{k} \right)\log\left( e_{ijk}\exp\left\{ \theta_{k} \right\}+\phi_{k} \right)$$

$$-\log\left\{ 1+\exp\left( -2\gamma_{k}-2\psi_{k}\sum_{\left| i^{'}-i \right|+\left| j^{'}-j \right|=1} z_{i^{'}j^{'}k} \right)\frac{\alpha_{\left\{ z_{ij1},\ldots-1,\ldots,z_{ijK} \right\}}}{\alpha_{\left\{ z_{ij1},\ldots1,\ldots,z_{ijK} \right\}}} \right\}.$$

$$\log P\left( z_{ijk}=-1 | \theta_{k},\phi_{k},\psi_{k},\gamma_{k},x_{ijk},e_{ijk} \right)=-\left( x_{ijk}+\phi_{k} \right)\log\left( e_{ijk}+\phi_{k} \right)$$

$$-\log\left\{ 1+\exp\left( 2\gamma_{k}+2\psi_{k}\sum_{\left| i^{'}-i \right|+\left| j^{'}-j \right|=1} z_{i^{'}j^{'}k} \right)\frac{\alpha_{\left\{ z_{ij1},\ldots1,\ldots,z_{ijK} \right\}}}{\alpha_{\left\{ z_{ij1},\ldots-1,\ldots,z_{ijK} \right\}}} \right\}.$$

Consider a special case where $K=2$, we hope to calculate the probability mass function for: $P(z_{ij1},z_{ij2}|\{\theta_{k}\},{\{\phi}_{k}\},\{\psi_{k}\},\{\gamma_{k}\}, {\{x}_{ijk}\},{\{e}_{ijk}\})$. Specifically, for a fixed $(i,j)$ pair, we have

$$\log P\left( z_{ij1},z_{ij2} | {\{z}_{-\left( ij1,ij2 \right)}\},\{\theta_{k}\},{\{\phi}_{k}\},\{\psi_{k}\},\{\gamma_{k}\}, {\{x}_{ijk}\},{\{e}_{ijk}\} \right)=Constant+\sum_{k=1}^{2} \left\{ \log\Gamma\left( x_{ijk}+\phi_{k} \right)-\log\Gamma\left( \phi_{k} \right)+x_{ijk}\left( \log e_{ijk}+\theta_{k}\frac{z_{ijk}+1}{2} \right)+\phi_{k}\log\phi_{k}-\left( x_{ijk}+\phi_{k} \right)\log\left( e_{ijk}\exp\left\{ \theta_{k}\frac{z_{ijk}+1}{2} \right\}+\phi_{k} \right) \right\}-\sum_{k=1}^{2} \log\left\{ 1+\frac{A\left( -z_{ijk} \right)}{A\left( z_{ijk} \right)} \right\}$$

Denoted as $B(z_{ij1},z_{ij2})$.

Therefore,

$$P(z_{ij1}=1,z_{ij2}=1|\{\theta_{k}\},{\{\phi}_{k}\},\{\psi_{k}\},\{\gamma_{k}\}, {\{x}_{ijk}\},{\{e}_{ijk}\})\propto\exp\left\{ B(1,1) \right\}$$

Considering all 4 possibilities:

$$P\left( z_{ij1}=1,z_{ij2}=1 | \{\theta_{k}\},{\{\phi}_{k}\},\{\psi_{k}\},\{\gamma_{k}\}, {\{x}_{ijk}\},{\{e}_{ijk}\} \right)=\frac{\exp\left\{ B(1,1) \right\}}{\exp\left\{ B(1,1) \right\}+\exp\left\{ B(1,-1) \right\}+\exp\left\{ B(-1,1) \right\}+\exp\left\{ B(-1,-1) \right\}}$$

Similarly, we can calculate $P\left( z_{ij1}=1,z_{ij2}=-1 | \{\theta_{k}\},{\{\phi}_{k}\},\{\psi_{k}\},\{\gamma_{k}\}, {\{x}_{ijk}\},{\{e}_{ijk}\} \right)$, $P\left( z_{ij1}=-1,z_{ij2}=1 | \{\theta_{k}\},{\{\phi}_{k}\},\{\psi_{k}\},\{\gamma_{k}\}, {\{x}_{ijk}\},{\{e}_{ijk}\} \right)$ and $P\left( z_{ij1}=-1,z_{ij2}=-1 | \{\theta_{k}\},{\{\phi}_{k}\},\{\psi_{k}\},\{\gamma_{k}\}, {\{x}_{ijk}\},{\{e}_{ijk}\} \right)$. We use the Gibbs sampler to update all the other parameters ($\theta_{k}$, $\phi_{k}$, $\psi_{k}$, $\gamma_{k}$). The hyper-parameter $\boldsymbol{\alpha}$ can be estimated from empirical data.

**Section 3: Implementation details of MUNIn and uni-sample analysis**

Uni-sample analysis was implemented following Xu *et al*.,^3^ where the initial peak status was randomly assigned. For MUNIn analysis, both peak status and parameters of each cell type, i.e., ($\theta_{k}$, $\phi_{k}$, $\psi_{k}$, $\gamma_{k}$), were initialized according to results from uni-sample analysis. Specifically, we searched within the range of +20% and -20% of estimates from uni-sample analysis for $\theta_{k}$, $\phi_{k}$, $\psi_{k}$, $\gamma_{k}$, which was equivalent to uniform priors. The across-sample dependency parameter $\alpha$ was estimated based on uni-sample inference. Then, the peak status and parameters were updated following the procedures described in Supplementary Section 1, and 10,000 Gibbs sampling steps were performed.

**Section 4: Real data evaluation framework**

To avoid false positive calls at a close distance, we further filtered the MUNIn-called peaks by excluding the bin pairs less than 50 kb apart. ROC curve was applied to illustrate the performance of MUNIn and uni-sample analysis, to help avoid the bias of unbalanced peak calling number between two methods. For the ROC curve, we excluded bin pairs with corresponding posterior probabilities less than the 10^th^ percentile, since including bin pairs with very low posterior probabilities is not meaningful. We partitioned the TADs within each chromosome into three categories (shared, GM-specific and IMR-specific peaks) according to their prior probabilities of four types of peak status (shared peaks, GM-specific peaks, IMR-specific peaks and shared backgrounds). If the prior probability is highest in shared peaks or cell-type-specific peaks, then we assigned this TAD to the corresponding category. If the prior probability is highest in shared backgrounds, then we assigned this TAD to the category where its prior probability is the second highest. Each ROC curve was then plotted only including TADs in that category.

**Section 5.** **Additional performance evaluation**

In this study, we further evaluated several other performance aspects of MUNIn. In our simulation, with loss of generality, we only considered the case that $\gamma_{k}$ = 0, where the proportion of peak and background is the same. To assess the robustness of MUNIn to the simulation parameters, we simulated data with different $\gamma_{k}$ values, -0.02, -0.05, -0.2 and -0.4. In all four scenarios, we observed that MUNIn achieved lower error rate in peak calling in all the three simulated samples than uni-sample analysis, when there is moderate or high dependency among samples (**Figure S5**). Our results demonstrated that MUNIn is robust to the simulation parameters.

In addition, we evaluated the scalability of MUNIn for a moderate sample size. We performed a simulation study with five samples. For all the five samples, the overall error rate for the peaks identified by MUNIn is substantially lower than that of uni-sample analysis when there is moderate or high dependency among samples (**Figure S6**).

To assess the robustness of MUNIn to the TAD boundaries, we re-ran the real data analysis for Hi-C data from GM12878 and IMR90 cell lines^2^ at 10kb resolution using a sliding window approach, instead of focusing on shared TADs. Specifically, we divided the genome into 1Mb windows (core region) with 200kb flanking regions on each side of the window and applied both MUNIn and uni-sample analysis to call interactions within each window. Similar to the results focusing on shared TADs, we observed that MUNIn obtained more accurate results for both GM12878 and IMR90-specific peaks (Figure 5a and d), while its performance in shared peaks was comparable to uni-sample analysis (**Figure S13**).

We then examined the overlapping between the cell-type-specific interactions and promoters and enhancers.^4^ In genome wide, we identified 535,908 and 674,194 GM12878- and IMR90-specific interactions. 109,406 and 41,178 (20.4 and 6.1%) of them are overlapped with promoter regions, which are significantly higher than those of all bin pairs (p-value < 2.2e-16 for both samples). Similarly, 71,978 and 48,485 GM12878- and IMR90-specific interactions (13.4 and 7.2%) are overlapped with enhancer regions, which are also significantly higher than the genome background (p-value < 2.2e-16 for both samples). These results indicate that the overlap between the significant interactions and promotor/enhancer region are not coincidences.

To further compare MUNIn with uni-sample analysis, we applied MUNIn and uni-sample analysis on Hi-C datasets of mouse embryonic stem cells for both wild-type and after CTCF deletion^5^ at 10kb resolution, and compared the peaks identified by uni-sample analysis and MUNIn with those identified by HiCCUPS.^2^ First, we overlapped bin pairs detected from uni-sample analysis and MUNIn with the union of HiCCUPS loops from the wild-type and the CTCF-depleted sample. Then, we looked at wild-type-specific peaks identified by uni-sample analysis or by MUNIn among those overlapping bin pairs. A bin pair was called as a wild-type-specific peak in uni-sample analysis if it was called as a peak in the wild-type sample and called as a background in the CTCF-depleted sample. Similarly, a bin pair is called as a wild-type-specific peak in MUNIn-analysis if the configuration of being a peak in wild-type sample while being a non-peak in the CTCF-depleted sample has the highest posterior probability. We ran both methods on chromosome 1 and found that among the overlapping bin pairs, uni-sample analysis called 17 wild-type-specific peaks, while MUNIn called 10 wild-type-specific peaks. We used HiCCUPS wild-type-specific loops as the ground truth and defined HiCCUPS wild-type-specific loops by first taking loops called by HiCCUPS in the wild-type mESC sample, and then excluding those that were also called as loops in the CTCF-depleted mESC sample by HiCCUPS. Uni-sample analysis had one false positive out of the 17 wild-type specific peaks, while all the 10 wild-type-specific peaks called by MUNIn were also identified by HiCCUPS. **Figure S14** shows aggregate peaks plots for wild-type-specific peaks identified by uni-sample analysis and MUNIn. We can see that MUNIn better captured the wild-type-specific pattern in mESC Hi-C data.

We used the same mESC HiC data^5^ to further compare MUNIn with FitHiC.^1^ Specifically, we performed FitHiC peak calling on the wild-type data (i.e., without CTCF depletion). We performed MUNIn peak calling by jointly analyzing Hi-C data before and after CTCF depletion (by inputting them as two samples into MUNIn) and focused only on peaks called in the wild-type sample (regardless of the status in the sample after CTCF deletion). We therefore were able to compare MUNIn and FitHiC peak calls in the wild-type sample. Specifically, we took the same number of top peaks from each method and compared percent overlapping with HiCCUPS loops (treated as the truth). For MUNIn, we first identified peaks by their inferred peak status, and then ranked them by posterior probability of being a peak in the wild-type sample from largest to smallest; for FitHiC, we ranked the peaks by their FitHiC p-values from smallest to largest. We compared the top 1,000 to 5,000 peaks called by each method and found that MUNIn had a higher number of overlaps with HiCCUPS loops than FitHiC (**Figure S15**). We chose this example and focus on the wild-type sample because HiCCUPS on the previously generated GM12878 wild-type data is believed to rather accurately reflect wild-type peaks. That our MUNIn results showing better performance than FitHiC demonstrates the power of MUNIn to more powerfully reveal peaks by borrowing information from another sample. In this case, it is particularly interesting because our results suggest that we attain better power detecting wild-type peaks in the wild-type sample even by borrowing information from the CTCF-depleted sample.

Finally, we estimated the computational time of MUNIn. Uni-sample analysis was first implemented on four shared TADs of different size from GM12878 and IMR90 cell lines, which contain 50, 100, 150 and 200 bins, respectively, and then MUNIn was performed on the results uni-sample analysis. The running time of uni-sample analysis and MUNIn was summed as the total computational time of MUNIn. For each TAD, the procedure was executed for 10 times. The results showed that MUNIn takes ~2 and 31 minutes to perform peak calling in a TAD consisting of 50 and 200 bins, respectively (**Figure S16**). We also assess the computational time of MUNIn at different resolutions. MUNIn was implemented on four 2 MB TADs of 10, 20 and 40 kb resolution for 10 times. The results showed that MUNIn takes only ~5 minutes for a TAD of 40 kb resolution, while ~36 minutes for a TAD of 10 kb resolution (**Figure S17**).

**Table S1.** Major characteristics of the benchmarking datasets.

| **Dataset** | **Cell type** | **Data type** | **GEO accession number/Download URLs** | **Ref** |
| --- | --- | --- | --- | --- |
| Simulations | - | Source codes | https://github.com/yycunc/MUNIn | - |
| Dixon *et al.* (2015) | Human embryonic stem cells | Dilute Hi-C | GSE52457 | 6 |
| Rao *et al.* (2014) | GM12878  IMR90 | In situ Hi-C | GSE63525 | 2 |
| Jung *et al.* (2019) | GM12878  IMR90 | Promoter-capture Hi-C | GSE86189 | 7 |
| Kubo *et al.* (2021) | Mouse embryonic stem cells | In situ Hi-C | GSE94452 | 5 |

**Supplementary Figures**

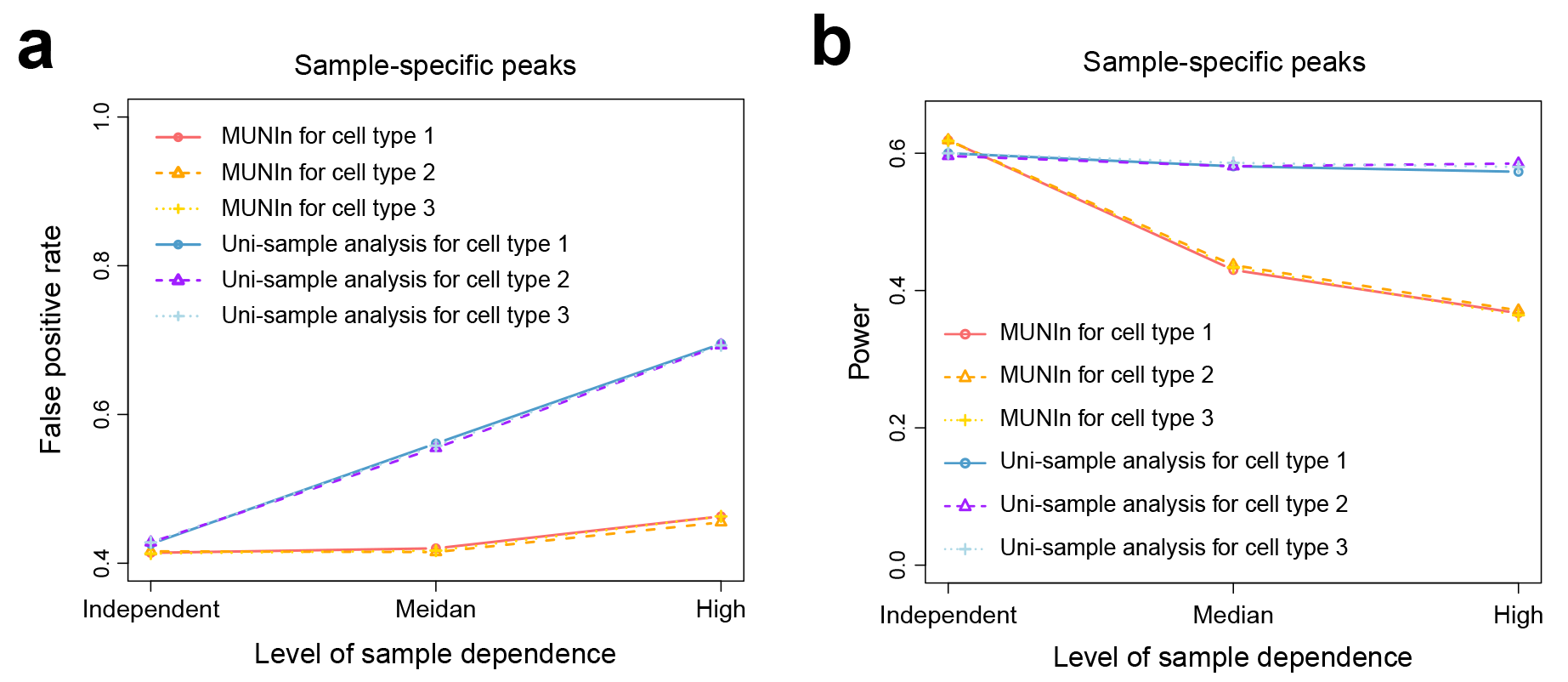

**Figure S1**. (**a**) False positive rate for the sample-specific peaks identified in each sample using MUNIn and uni-sample analysis in the simulated data under the scenario that all three samples have equal sequencing depth. (**b**) Power for the sample-specific peaks identified by MUNIn and uni-sample analysis.

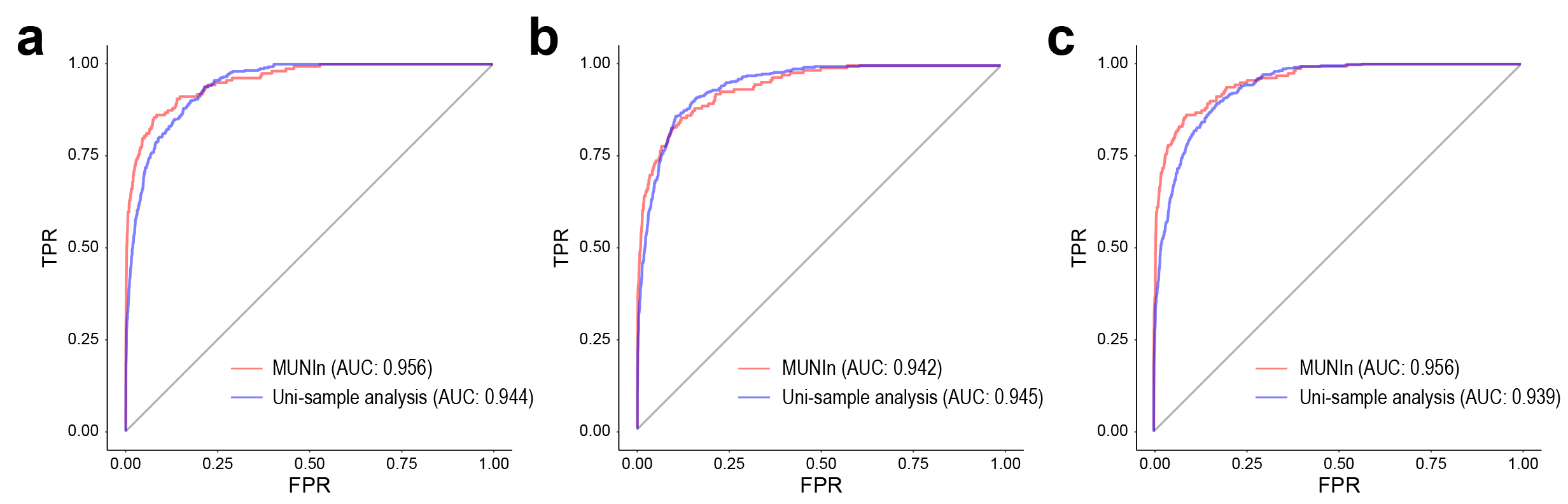

**Figure S2.** ROC curves for sample 1-, 2- and 3-specific peaks identified by MUNIn and uni-sample analysis in the simulated data under the scenario that all three samples have equal sequencing depth.

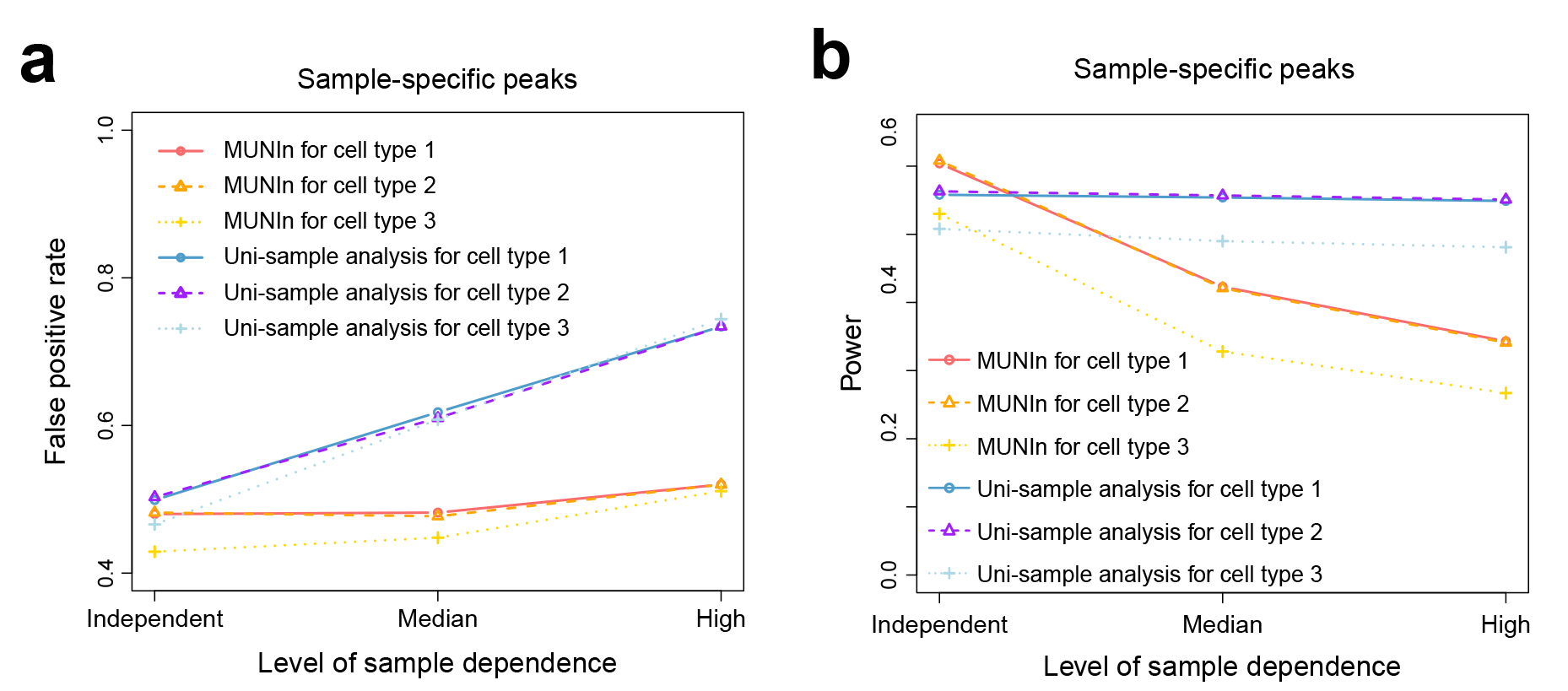

**Figure S3.** (**a**) False positive rate for the sample-specific peaks identified in each sample using MUNIn and uni-sample analysis in the simulated data under the scenario that the three samples have different sequencing depth. (**b**) Power for the sample-specific peaks identified by MUNIn and uni-sample analysis.

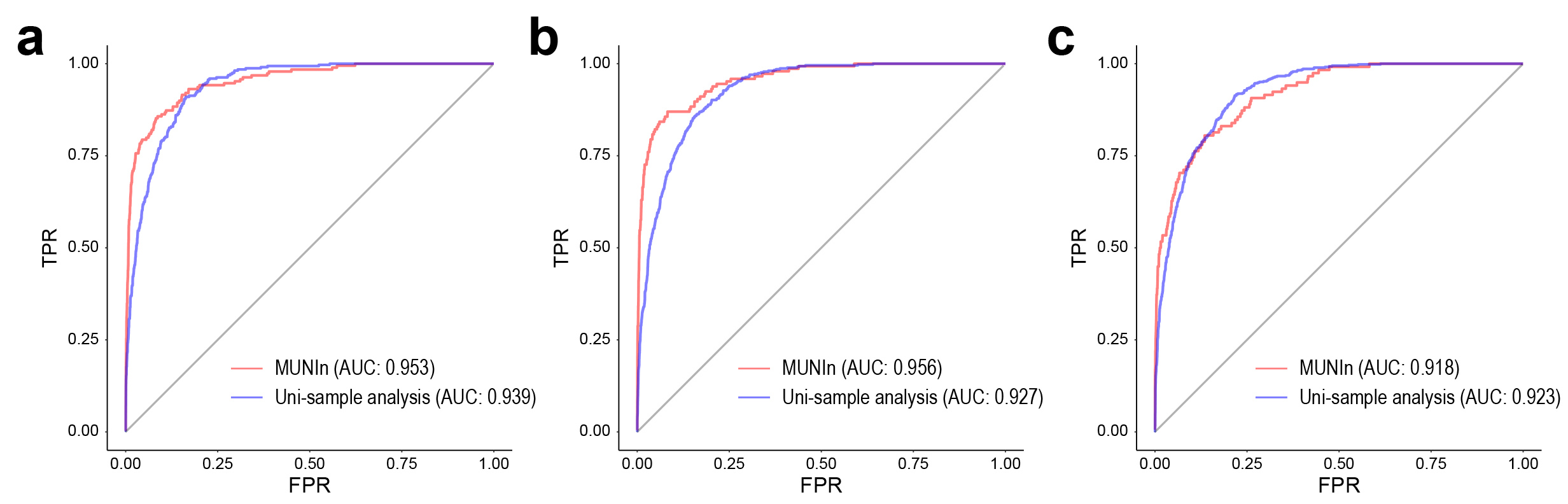

**Figure S4.** ROC curves for sample 1-, 2- and 3-specific peaks identified by MUNIn and uni-sample analysis in the simulated data under the scenario that the three samples have different sequencing depth.

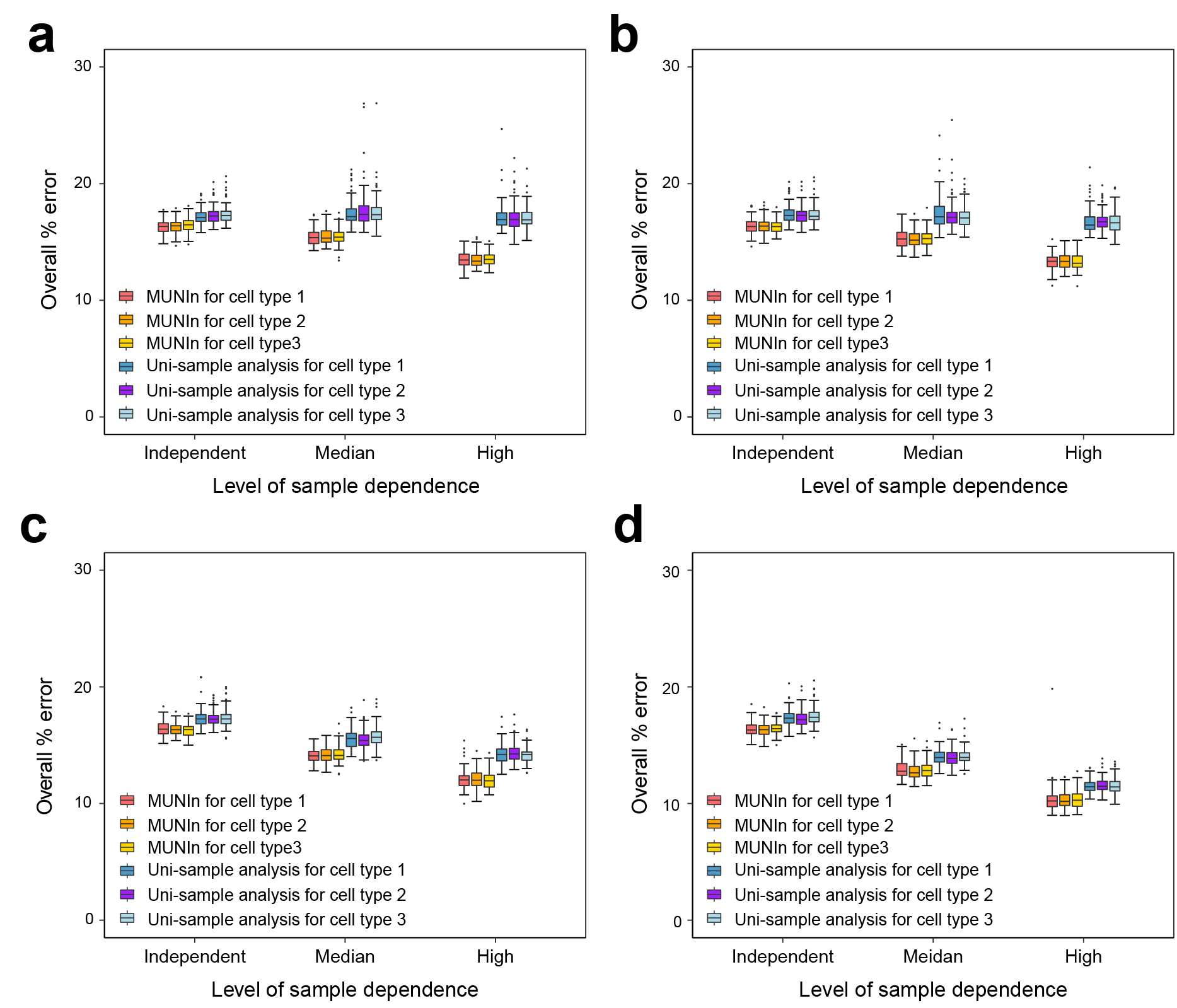

**Figure S5.** The overall error rate (denoted as “%error”) in peak identification in each sample of the simulations using different $\gamma_{k}$ values, (**a**) -0.02, (**b**) -0.05, (**c**) -0.2 and (**d**) -0.4.

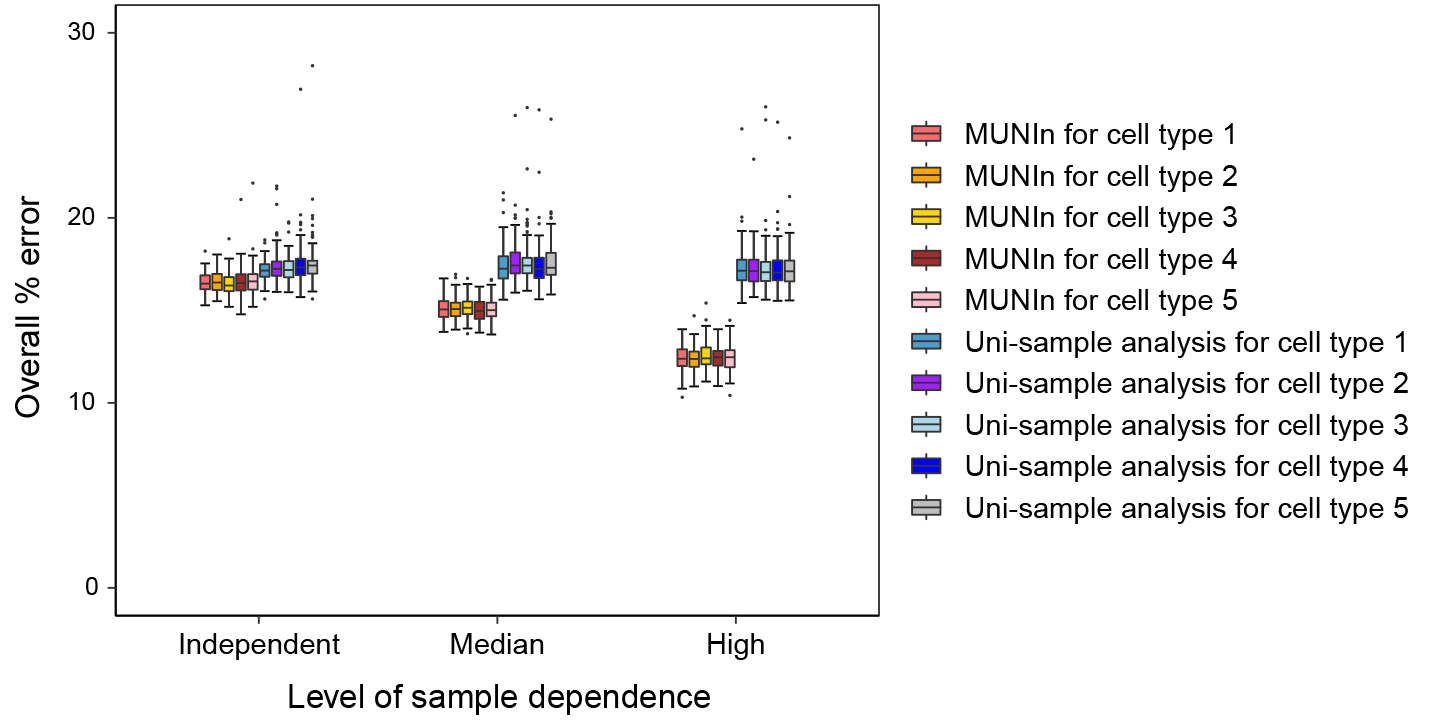

**Figure S6.** The overall error rate (denoted as “%error”) in peak identification in each of the five simulated samples using MUNIn and uni-sample analysis, respectively.

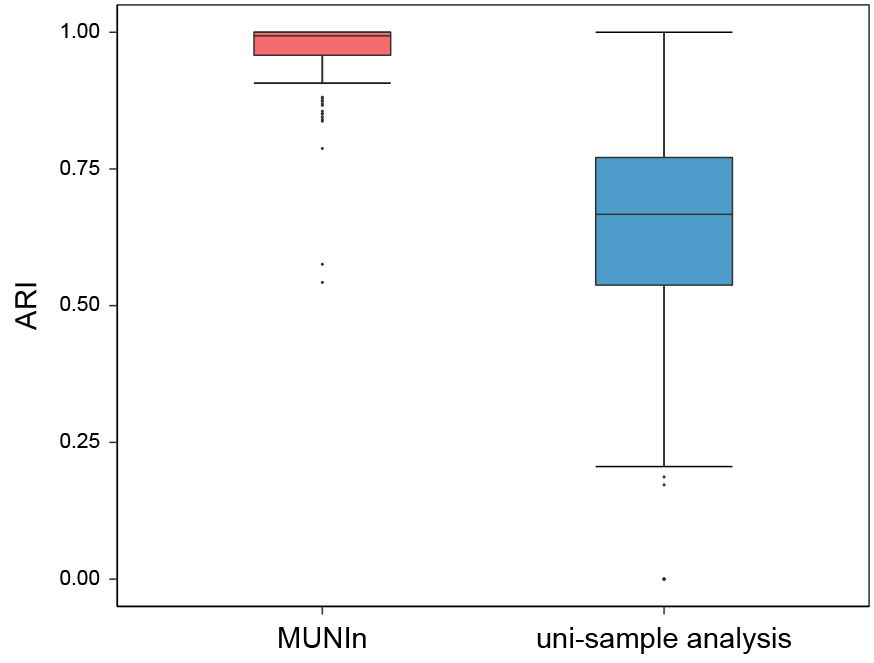

**Figure S7.** Adjusted Rand Index (ARI) showing the consistency between the interactions detected in the two biological replicates of human embryonic stem cells.

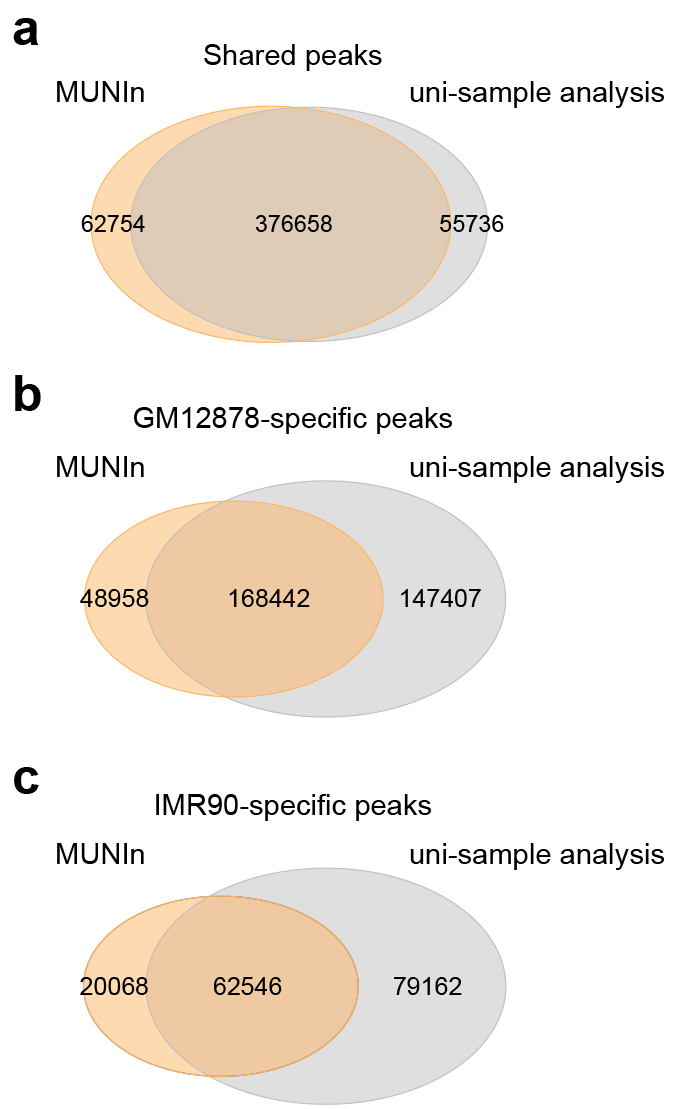

**Figure S8.** Venn diagram showing the overlap of the (**a**) shared, (**b**) GM12878-specific and (**c**) IMR90-specific peaks identified by MUNIN and uni-sample analysis.

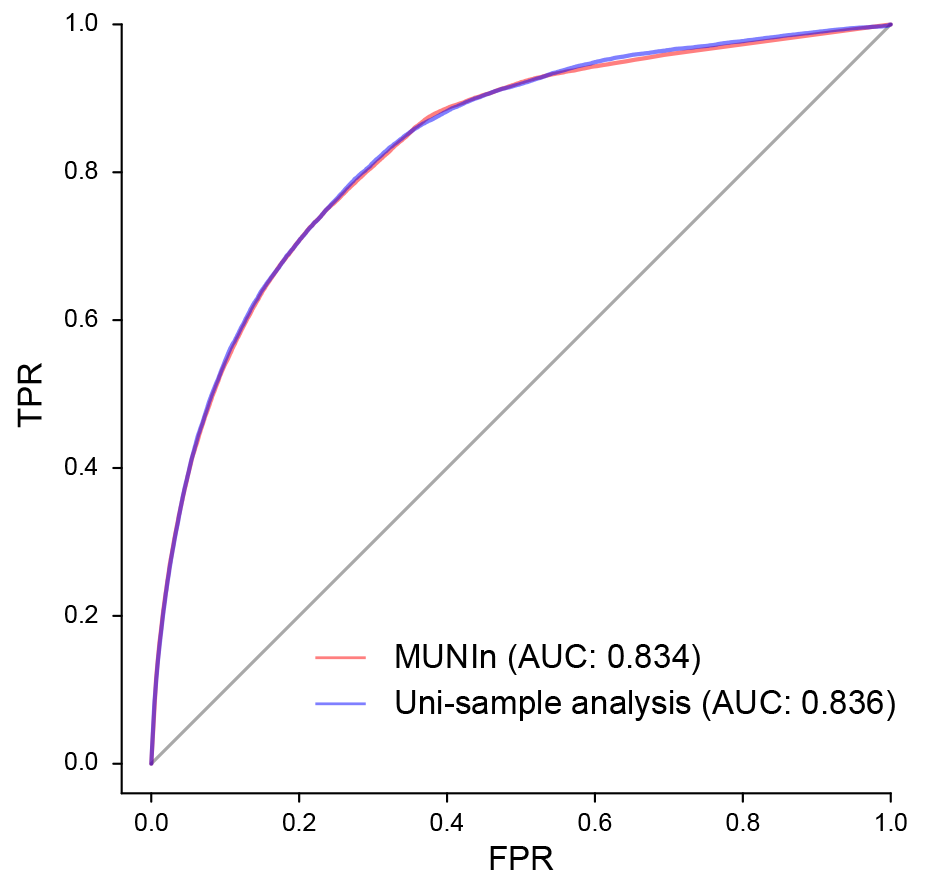

**Figure S9.** ROC curve for the shared peaks detected in both GM12878 and IMR90 using MUNIn and uni-sample analysis.

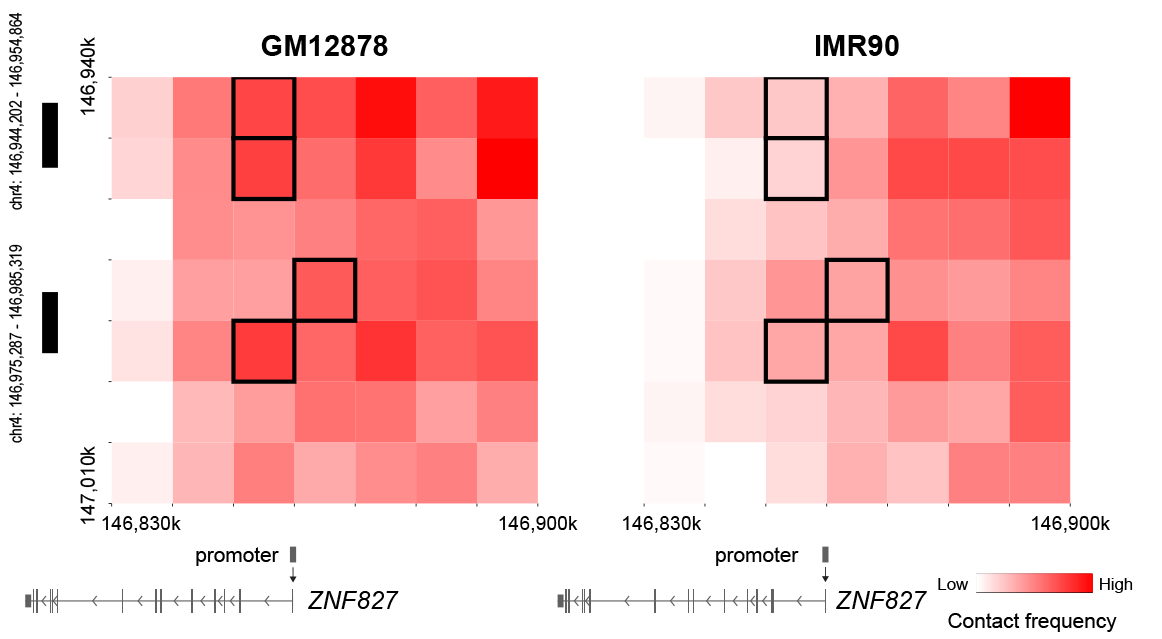

**Figure S10.** Heatmap showing the GM12878-specific peaks in GM12878 (left) and IMR90 (right) Hi-C data. One bin of these pairs (highlighted in black) is overlapped with the promoter of *ZNF827* gene (transcription start site (TSS) +/- 500bp), while the others are overlapped with known typical enhancers (chr4:146,975,287-146,985,319 and chr4:146,944,202-146,954,864) in GM12878 cells. Gene model is obtained from WashU epigenome browser.^8^

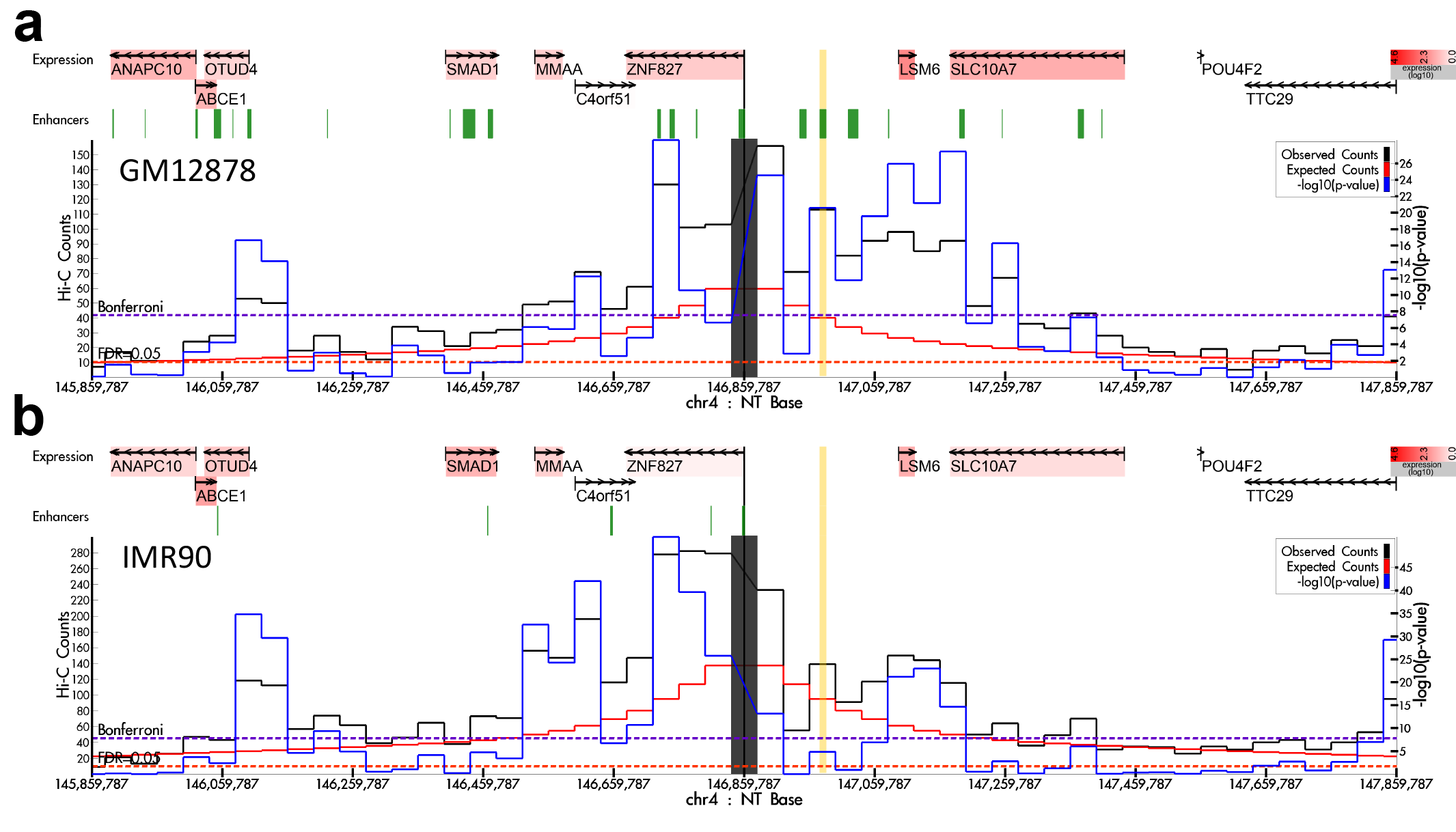

**Figure S11.** Virtual 4C plot showing one example of the GM12878-specific peaks in (a) GM12878 and (b) IMR90 Hi-C data using HUGIn.^9^ On the top of each panel, the genes are colored according to their expression level (the deeper red, the higher the expression level) with arrows indicating the direction of transcription and a vertical bar indicating the transcription start site (TSS). On the bottom of each panel, the chromatin interaction from Hi-C data is shown by a virtual 4C plot. The anchor bin overlapped with the promoter region of gene *ZNF827* is indicated as a thick grey vertical bar at the center. The bin overlapped with the GM12878-specific enhancer region is highlighted in yellow. The black line shows the observed counts; the red line shows the expected counts; and the blue line shows the -log10(p-value). The range of observed and expected counts is shown on the left Y-axis; and the range of the -log10(p-value) is plotted on the right Y-axis. The X-axis is the genomic location on chromosome 4.

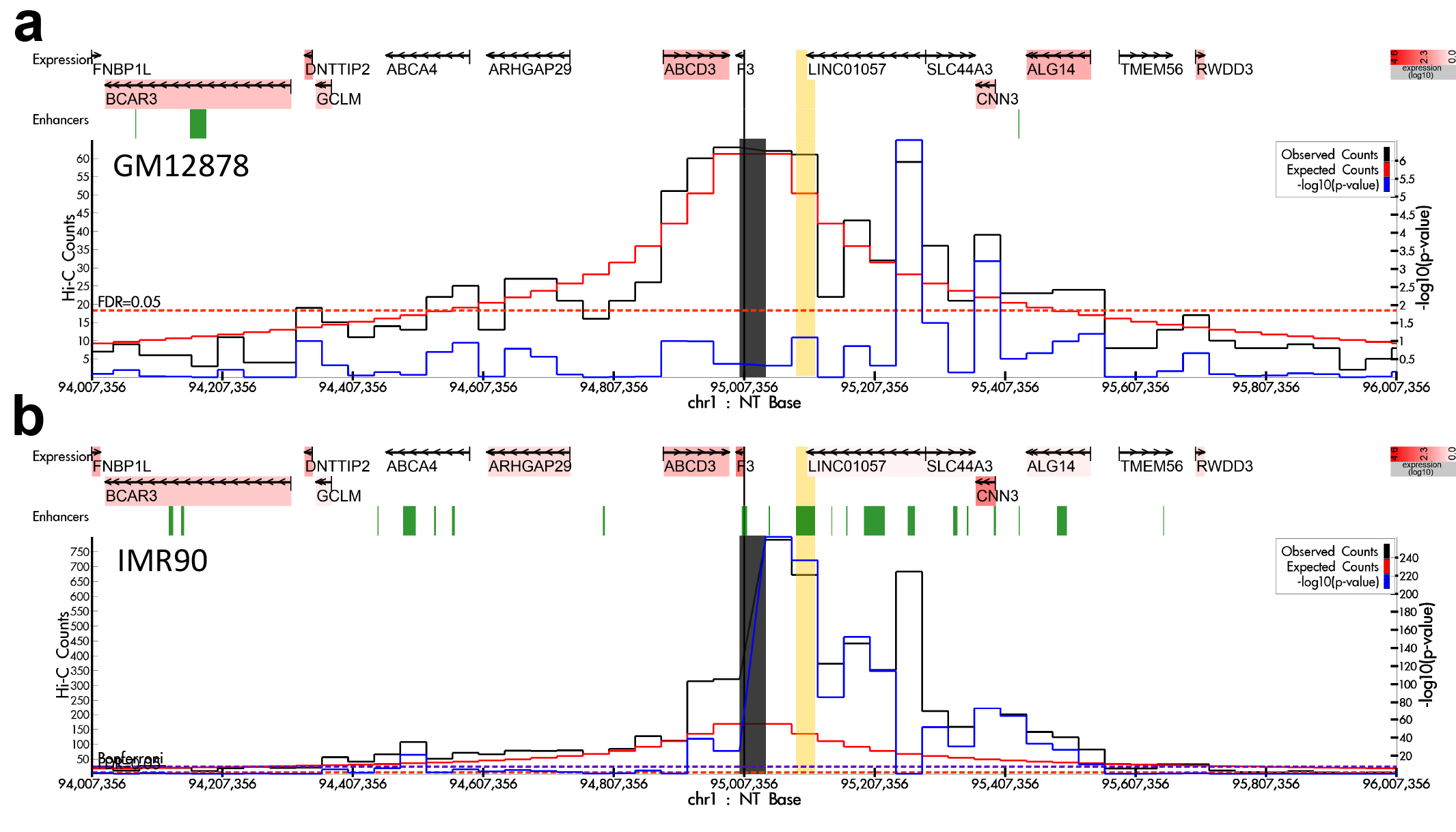

**Figure S12.** Virtual 4C plot showing one example of the IMR90-specific peaks in (a) GM12878 and (b) IMR90 Hi-C data using HUGIn.^9^ On the top of each panel, the genes are colored according to their expression level (the deeper red, the higher the expression level) with arrows indicating the direction of transcription and a vertical bar indicating the transcription start site (TSS). On the bottom of each panel, the chromatin interaction from Hi-C data is shown by a virtual 4C plot. The anchor bin overlapped with the promoter region of gene *F3* is indicated as a thick grey vertical bar at the center. The bin overlapped with the IMR90-specific enhancer region is highlighted in yellow. The black line shows the observed counts; the red line shows the expected counts; and the blue line shows the -log10(p-value). The range of observed and expected counts is shown on the left Y-axis; and the range of the -log10(p-value) is plotted on the right Y-axis. The X-axis is the genomic location on chromosome 1.

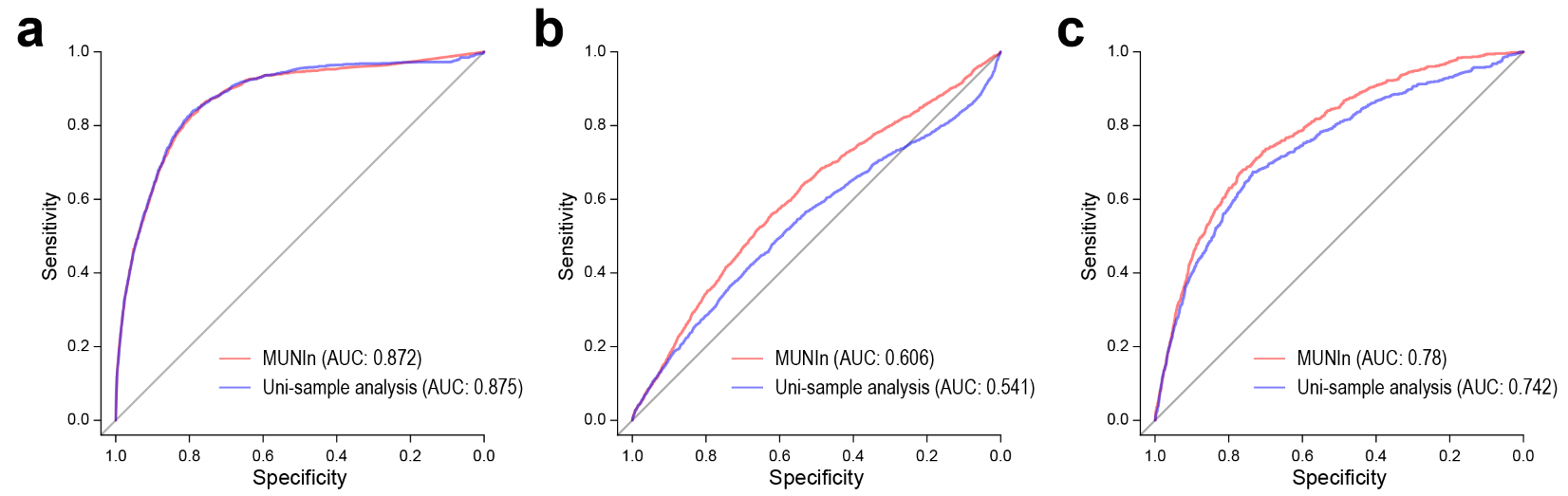

**Figure S13.** ROC curves for (**a**) shared, (**b**) GM12878-specfic and (**c**) IMR90-sepcific peaks identified by MUNIn and uni-sample analysis in the sliding window approach.

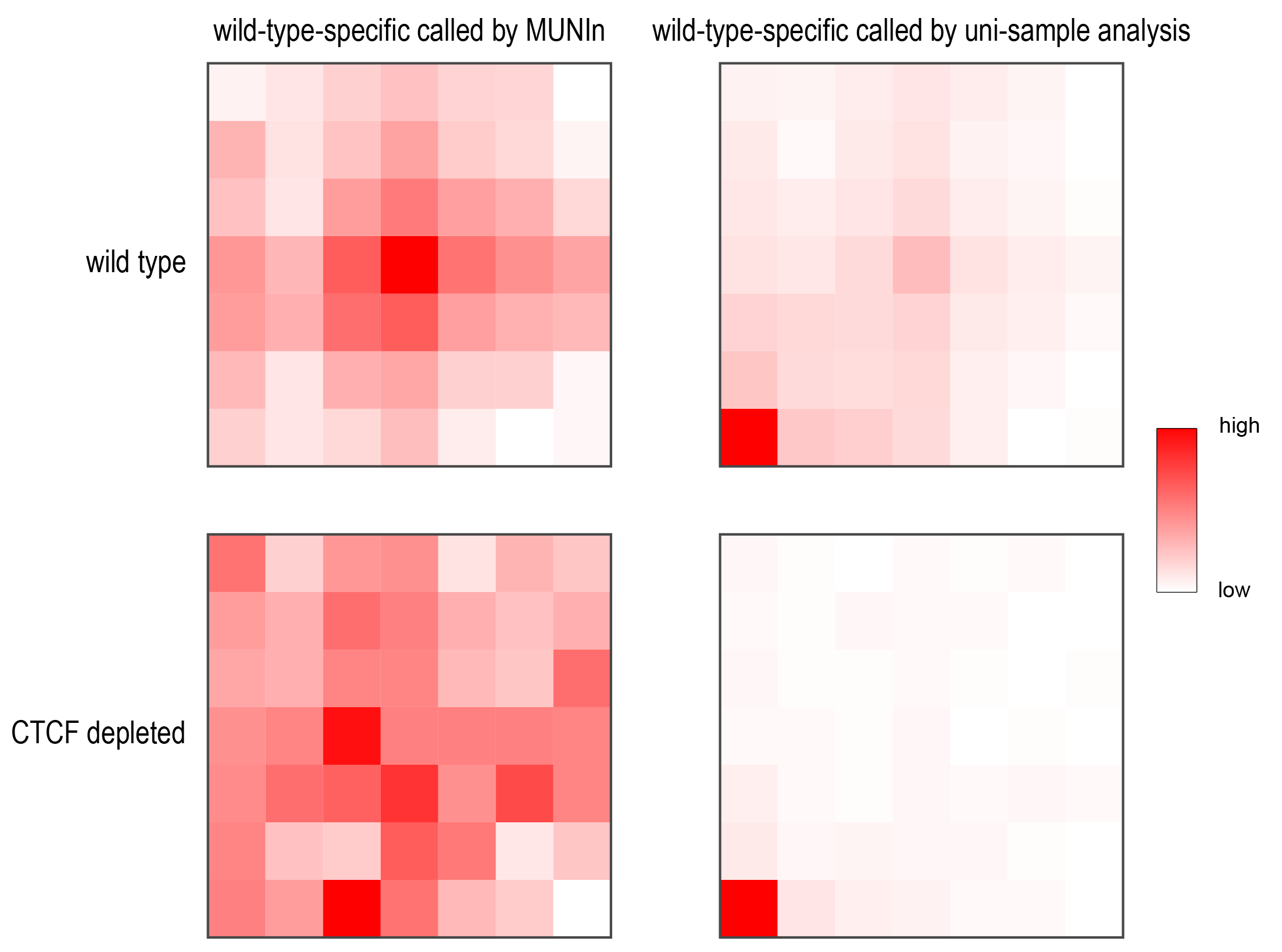

**Figure S14.** Aggregate peaks plots on mESC wild-type-specific peak loci identified by MUNIn and uni-sample analysis. Observed contact counts are aggregated over the peak bin pairs and their 7 x 7 neighbors. Each bin has length 10kb. The first row shows aggregate counts for the wild-type sample and the second row shows aggregate counts for the CTCF-depleted sample. For wild-type-specific peaks, we expect to see a peak pattern in the wild-type sample but not in the CTCF-depleted sample**.**

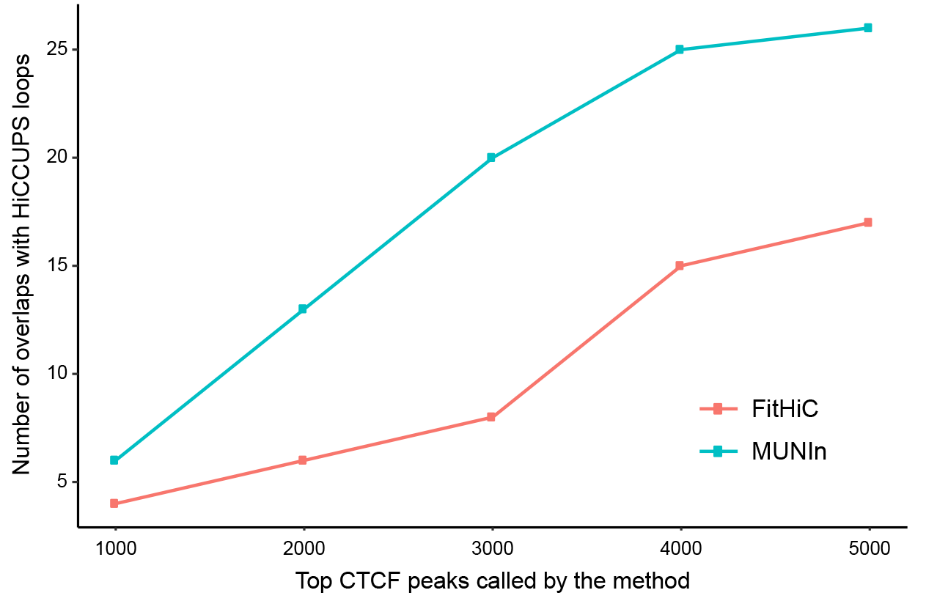

**Figure S15.** Number of peaks overlapped with HiCCUPS loops at the top 1000, 2000, 3000, 4000 and 5000 CTCF peaks called by MUNIn and Fithic.

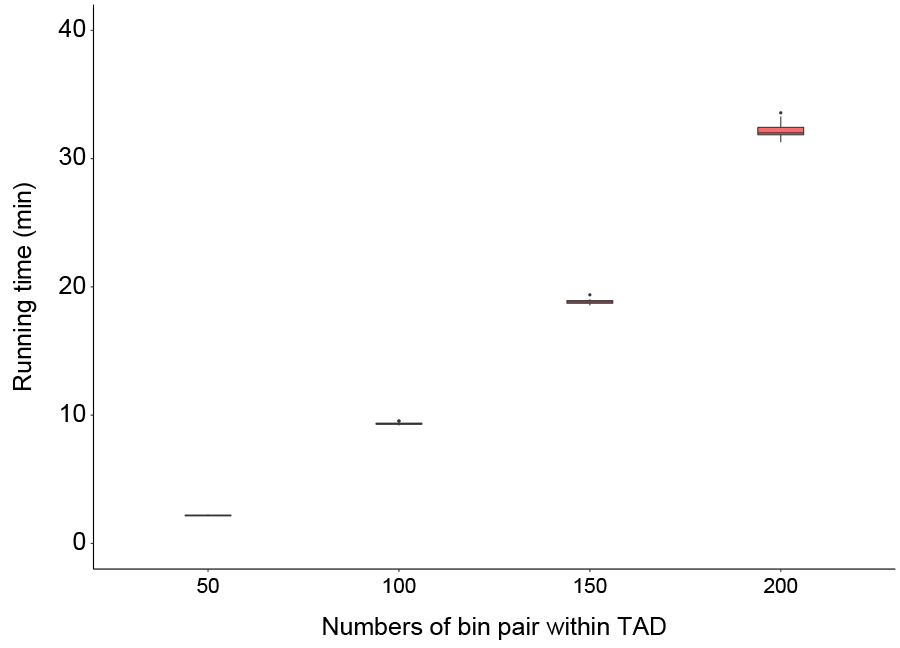

**Figure S16.** Running time of MUNIn analysis for four shared TADs of GM12878 and IMR90 cell lines, which contain 50, 100, 150 and 200 10kb bins, respectively. For each TAD, uni-sample analysis and MUNIn were executed 10 times. Running time (Y-axis) is in minutes (min). The computing time is from running on a Linux-based computing cluster.

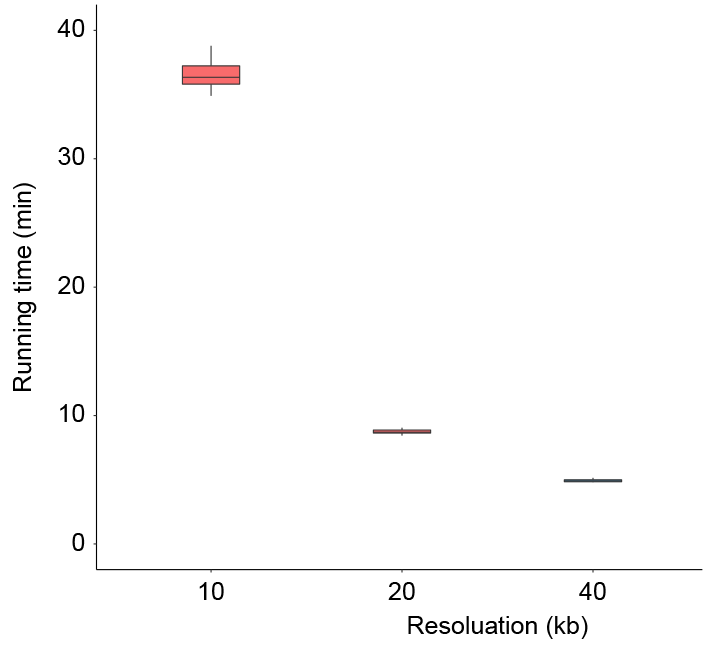

**Figure S17.** Running time of MUNIn analysis for four shared TADs of GM12878 and IMR90 cell lines at different resolutions, 10, 20 and 40kb. Running time (Y-axis) is in minutes (min). The computing time is from running on a Linux-based computing cluster.
